# Noradrenergic cross-modular reciprocal inhibition within the locus coeruleus

**DOI:** 10.1101/2022.09.07.506929

**Authors:** Oscar Davy, Ray Perrins, Marina Lavigne, Eric Kremer, Krasimira Tsaneva-Atanasova, Michael Ashby, Anthony E Pickering

## Abstract

The Locus Coeruleus (LC) is the primary noradrenergic nucleus in the brain with widespread projections driving changes in cognitive state and animal behaviour. The LC is composed of multiple “modules” with specific efferent target domains enabling discretional neuromodulation. LC neuronal activity releases noradrenaline within the nucleus as a local feedback mechanism, but it is not known how this influences modular output. We address this question using whole-cell recordings and calcium imaging in rat pontine slices in combination with LC neuronal ensemble modelling to assess the influence of local noradrenaline release on cross-modular interactions.

Electrophysiological recordings of LC neurons from rats transduced with the optogenetic actuator ChR2 showed auto-inhibition and lateral inhibition (of surrounding non-transduced neurons). This inhibition was strongly frequency dependent and was mediated by noradrenaline acting on alpha2-adrenceptors (α2R). To allow calcium-imaging of LC neuronal ensembles a Canine-Adenoviral vector strategy was developed using the PRS promoter to drive selective expression GCaMP6s. Calcium imaging allowed resolution of both increases and decreases in LC activity (to TTX / clonidine or high potassium). Selective chemogenetic-activation of subsets of LC neurons (expressing the ionotropic actuator PSAM) revealed both a direct excitation (after application of PSEM308, 3-30µM) and an α2R-mediated inhibition of neighbouring LC cells (non-transduced). Differential retrograde targeting of PSAM or GCaMP6s to specific LC modules showed the presence of strong, reciprocal cross-modular inhibition (shown for the LC-olfactory bulb vs LC bulbospinal modules) and a subsequent rebound activity inversion.

This represents a preferential, targeted, cross-modular, lateral inhibition within the LC rather than a non-specific surround inhibition. Computational modelling showed the emergence of lateral inhibition and biphasic responses to modular activation when α2R signalling and noradrenergic reuptake saturation were included. This interaction may facilitate recruitment of neuronal ensembles by coherent inputs and represents a bottom-up differential contrast-enhancement mechanism within the LC to produce a modality specific focus.

## Introduction

The Locus Coeruleus (LC) is the primary source of noradrenaline (Dahlstroem & Fuxe, 1964), consisting of bilateral, densely-packed cell clusters in the pons, with approximately 3,000 neurons in rats and 50,000 in humans (Swanson, 1976; Sharma *et al*., 2010). LC neurons extend remarkable axonal projection trees to much of the brain to release noradrenaline. In behaving animals, the LC neurons display tonic ‘pacemaker’ firing at rates of ∼0.5-5Hz, with the rate of this tonic activity broadly correlating with levels of alertness (Aston-Jones & Bloom, 1981; Aston-Jones & Cohen, 2005) and this activity ceases during rapid eye movement (REM) sleep (Hayat *et al*., 2020; Kjaerby *et al*., 2022). In addition to ongoing tonic activity, LC neurons also display a burst-firing mode of activity, known as phasic, in response to novel or emotionally salient stimuli. This phasic firing has been suggested to signal internal and external state changes that are important for cognition and learning (Berridge & Waterhouse, 2003; Aston-Jones & Cohen, 2005; Sales *et al*., 2019).

In line with its extensive reach, the LC is implicated in a wide range of brain functions in health including salience detection and sensory discrimination, memory processing and learning, cognitive flexibility, motor control and motivation, arousal and sleep-wake transitions (Samuels & Szabadi, 2008; Sara, 2009; Aston-Jones & Waterhouse, 2016). Disorders of noradrenergic signalling and LC function have also been implicated in multiple common brain pathologies such as Alzheimer’s disease, Parkinson’s disease, depression, anxiety, post-traumatic stress disorder and chronic pain (Benarroch, 2009; Llorca-Torralba *et al*., 2016; Weinshenker, 2018).

These diverse influences across multiple domains of the neuroaxis with specificity in functional roles have motivated many studies to define the underlying cellular / network architecture and adrenergic receptor expression pattens that enables these selective noradrenergic neuromodulatory effects. Multiple lines of evidence now indicate that the LC is organised into discrete modules with each neuron having a preferential axonal projection distribution (reviewed in (Foote *et al*., 1983; Chandler *et al*., 2019; Poe *et al*., 2020)). This organisation potentially allows the LC to provide discrete neuromodulation of target fields rather than simply providing an obligatory global noradrenergic signal. The advent of genetic circuit manipulation techniques capable of dissecting the influence of this modularity has allowed dissociation of specific noradrenergic actions mediated by distinct projections to be identified such as: pain suppression vs aversion (Hirschberg *et al*., 2017), fear memory embedding vs extinction (Uematsu *et al*., 2017) and motor execution vs negative reinforcement learning (Breton-Provencher *et al*., 2022).

This modular LC architecture with clusters of cell somata, projecting to different areas of the neuroaxis, but grouped tightly together within the pons raises questions about how the specificity of input-output relationships is mapped and preserved. At a cellular level there is a prevalent view of the LC as a relatively homogenous group of catecholaminergic neurons with similar electrophysiological properties exhibiting pacemaker activity (Williams *et al*., 1984; Williams & Marshall, 1987) and with many commonalities in terms of their synaptic inputs (Schwarz *et al*., 2015; Breton-Provencher *et al*., 2022), genetic profile and functional responses. There is also evidence for gap junctional coupling of the spontaneous activity within the LC that is strongest in the developing animal (Williams *et al*., 1984; Williams & Marshall, 1987; Christie *et al*., 1989; Ishimatsu & Williams, 1996; Alvarez *et al*., 2002). That picture of homogenous, synchronous and uniform activity has been challenged by a number of *in vitro* and *in vivo* studies which have shown distinct cellular properties (Chandler *et al*., 2014; Li *et al*., 2016; Totah *et al*., 2018; Wagner-Altendorf *et al*., 2019), developmental pedigree (Plummer *et al*., 2017) and a lack of synchrony between cell pairs (Totah *et al*., 2018). Nonetheless the details of how these differences in properties enable discrete actions of LC modules to be enacted is currently lacking.

A distinctive feature of the LC neurons is a feedback or lateral inhibition mediated by noradrenaline released within the nucleus (Callado & Stamford, 1999; Pudovkina *et al*., 2001) and acting via α2Rs-adrenoceptors which are expressed at high density (Tavares *et al*., 1996) to produce a powerful inhibition of LC activity (Aghajanian *et al*., 1977; Ennis & Aston-Jones, 1986). This is mediated by activation of G-protein coupled inward rectifying potassium channels (GIRKs) (Williams *et al*., 1985). This has typically been characterised as a braking mechanism to restore activity to basal levels after an activation such as that seen after a noxious stimulus with a phasic burst of activity followed by a period of suppressed spontaneous discharge that gradually returns to baseline (Cedarbaum & Aghajanian, 1978; Ennis & Aston-Jones, 1986). It has also been noted that there is a tonic level of noradrenaline release within the nucleus such that spontaneous LC discharge is also under a degree of ongoing α2R-mediated inhibition (Grandoso *et al*., 2004). A biological role for this powerful feedback mechanism beyond a braking action has not been convincingly demonstrated although it has recently been suggested, on the basis of modelling, that “bystander inhibition” may influence the output of the LC whereby active neurons might be able to influence their neighbours by the local release and diffusion of noradrenaline (Baral *et al*., 2022). We have investigated this possible mechanism in the context of networks and modules of LC neurons to show that cross-modular inhibition powerfully regulates the excitability of LC ensembles and that this can act as a bottom-up contrast boosting mechanism to alter the output of opposing LC modules.

## Materials & Methods

All animal experiments were undertaken in accordance with the UK (Animals) Scientific Procedures Act 1998 and approved by the University of Bristol Animal Welfare and Ethical Review Board. Animals were housed in an enriched environment, under a reversed 12:12 lighting regimen, with food and water freely available.

### Stereotactic vector injections

P21-28 male Han-Wistar rats (Animal Services Unit breeding colony) were anaesthetised by intraperitoneal (I.P.) injection of ketamine (50mg/kg) & medetomidine (300μg/kg). Animals were head-fixed in a stereotaxic frame (Neurostar Drill and Injection robot with a Kopf frame), with a rectal thermometer and homeothermic heat blanket, maintained at 37 °C. Under aseptic conditions, the skull was exposed, and right sided burr hole was drilled for each vector injection (co-ordinates in Table 1). An injection pipette made from pulled capillary glass (Drummond Scientific, 1mm diameter) was filled with mineral oil and back filled with viral vector. The pipette was inserted to the target depth and viral vectors were injected at a rate of 100nL/minute, (using the titres shown in Table 2). The only adaptation was for the retrograde targeting of bulbo-spinal LC efferents. For these injections the dorsal medulla was accessed via the atlanto-occipital membrane (Cerritelli et al., 2016). The rat was head-fixed in a stereotaxic frame using ear-bars with the bite bar placed on the dorsal surface of the snout, and adjusted so the nose pointed downwards at ∼30°. A skin incision was made on the caudal aspect of the scalp, following the midline, and the muscle underneath was blunt dissected to reveal the atlanto-occipital membrane. A hypodermic needle was used to puncture the atlanto-occipital membrane, and vector injections were made unilaterally (see Table 1, AP coordinates relative to the caudal tip of the area postrema i.e. calamus scriptorius). Following vector injection and pipette withdrawal, the muscle tissue was sutured, followed by the skin incision. After injection at the final site in each tract, the pipette was left in place for 5 minutes to ensure effective vector dispersion into the tissue. Following surgical closure, animals were given atipamezole (1mg/kg, i.p.) to reverse the anaesthesia, as well as 1ml saline (sub cutaneous, s.c.) and buprenorphine (30μg/kg, s.c.), and allowed to recover on a heat-mat.

**Table 1.** Stereotaxic Injection Coordinates.

|  | *Antero-Posterior* | *Medio-lateral* | *Depth\** | *Angle* |
| --- | --- | --- | --- | --- |
| *LC* | -1<br>(vs *lambda*) | +1.1 | 4.6, 4.9, 5.2, 5.5 | 10° rostral |
| *Olfactory Bulb* | +6.5, +7.5<br>(vs *Bregma*) | 0.8 | 1, 1.5, 2 | 0° |
| *Bulbo-spinal* | -1.5<br>(from C.S **) | 1.5 | 0.25, 0.5, 0.75, 1 | 0° |

*All measurements are given in mm*
*\* Injections made at these depths from brain surface for all injection tracts*
*\*\* C.S. - Calamus Scriptorius, see methods section*

**Table 2.** Viral Vectors.

| *Vector* | *Source* | *Titre* |
| --- | --- | --- |
| *CAV2-PRS-GCaMP6s* | *Eric Kremer, IGMM\** | *1.3x10<sup>11</sup> p.p./ml* |
| *CAV2-PRS-PSAM-eGFP* | *Eric Kremer, IGMM\** | *2.0x10<sup>11</sup> T.U./ml* |
| *CAV2-PRS-ChR2-mCherry* | *Eric Kremer, IGMM\** | *1.2x10<sup>11</sup> T.U./ml* |

*\* Institut de Génétique Moléculaire de Montpellier, Montpellier, France; Université de Montpellier*

### *Ex-vivo* slice preparation

After 14-21 days recovery from viral vector injection, pontine slices were prepared for whole cell recordings using methods as previously described (Howorth *et al*., 2009a; Hickey *et al*., 2014; Li *et al*., 2016). In brief, rats were terminally anaesthetised with isoflurane (4%) and decapitated. The brain was removed from the cranium, placed ventral side down onto filter paper in an ice-cold glass petri-dish, and rapidly dissected using a razor blade to make transverse cuts rostrally and caudally to produce a pontine tissue block. The rostral face of the tissue block was glued onto the cutting stage of a vibrating microtome (DSK Pro-linear slicer, Japan) placed into ice-cold carbogenated cutting aCSF (in mM: sucrose 58.4, NaCl 85, KCl 2.5, NaHCO_3_ 26, NaH_2_PO_4_ 1.25, MgCl_2_ 2, CaCl_2_ 2, D-Glucose 10, 290mOsm/L osmolality). Transverse slices were cut (300μm) using the vibrating microtome, and allowed to recover at 37°C in carbogenated recording aCSF (same as cutting aCSF, but with NaCl substituted for sucrose, to a final concentration of 126mM) for at least 1 hour. Slices were then kept at room temperature (21°C) carbogenated recording aCSF prior to recordings.

### Whole-cell recordings

Whole-cell patch clamp recordings were made from LC neurons. Slices were placed in a chamber on an upright fluorescence microscope (DMLFSA, Leica Microsystems) and cells were visualised with gradient contrast illumination. Carbogenated aCSF was perfused at 1ml/minute from a peristaltic pump with separate reservoirs for aCSF±drug application. The aCSF was warmed to 33°C by an in-line heater and temperature controller (Warner Instruments SH-27B & TC-324C). Borosilicate glass electrodes (GC120F-10, Harvard Apparatus) were pulled to ∼3MΩ tip resistance using a pipette puller (P-87, Sutter Instrument Co.). Pipettes were filled with internal solution (in mM: K Gluconate (130), KCl (10), Na-HEPES (10), MgATP (4), EGTA (0·2) and Na_2_GTP (0·3)). Neurons were selected based on their location within the LC nucleus - visible as a relatively translucent area in the lateral aspect of the floor of the fourth ventricle and confirmed during recording by their characteristic electrophysiological profile with slow (<10Hz) tonic firing rates. Viral vector transduced LC neurons (expressing ChR2) were identified by mCherry fluorescence, visualised under epifluorescent illumination (Chroma filterset 41034). Light for optogenetic stimulation from a 473nm LED source (11mW, Doric Lenses) was routed via an optical fibre (400μM diameter) positioned above the LC.

Whole-cell recordings were made by obtaining a gigaohm seal on the plasma membrane and applying brief negative pressure using a glass syringe in order to gain access. The electrical signals were recorded using a Multiclamp 700a amplifier (Axon Instruments) in current / voltage clamp modes. Only recordings in which action potentials remained stable (overshooting zero), and in which resting membrane potential remained below −40mV during a basal period of 300 seconds included for analysis. Data was low pass filtered (3KHz) and digitized at 10KHz by a Power1401 AD converter and displayed / stored on a pc using the Spike2 acquisition software (v7.14, Cambridge Electronic Design).

Measurement of action potential and membrane oscillation parameters was performed in Spike2 (CED) using custom written scripts. All action potential parameters were averaged across 20 action potentials per cell, then across all cells in the dataset. The threshold potential for sodium-based spiking was defined as the membrane potential at which the change in membrane voltage exceeded 7.5V/sec. For Calcium-based action potentials (in the presence of TTX), the threshold was defined as the membrane potential at which the change in membrane voltage exceeded 2 V/sec - due to the slower temporal dynamics of the upstroke of calcium action potentials. Action potential amplitude was measured from the threshold potential to the peak potential for each spike. Action potential duration was measured as the time between the voltage trace passing 1/3 of the amplitude and returning to 1/3 of the amplitude for each spike.

Statistical analysis was performed using parametric and non-parametric tests (as appropriate) implemented in Python3, using the Scipy Stats module ((Virtanen *et al*., 2020), and aggregate plots generated in Python3. Results are reported as mean±standard deviation, unless otherwise stated.

### 2-photon imaging

Slices were placed in the recording chamber of a 2-photon microscope (Bruker/Prairie Technologies Ultima IV), supplied constantly with carbogen bubbled aCSF, warmed to 33°C with an in-line heater and temperature controller (Warner Instruments SH-27B & TC-324C), with a 5ml bubble-trapping reservoir for the aCSF feed, as well as a reservoir for aCSF+drug applications. Illumination from a 470nm LED (Thorlabs) with a 550nm bandpass emission filter was used to visually identify GCaMP6s positive cells and orient the objective lens (Nikon MRP07220, 16x magnification, 0.8 numerical aperture) above the transduced LC.

The microscope was switched to 2-photon mode, and excitation light (920nm for GCaMP6s excitation) was raster scanned across the field of view (Spectra-Physics Mai-Tai laser). Emitted light was collected by the objective lens and routed via a 565nm dichroic mirror (Chroma T565lpxr) to a GaAsP photocathode photomultiplier tube (Hamamatsu) with an emission bandpass filter at 525/70nm. The 2-photon recordings were typically captured at ∼1Hz imaging frequency using PrairieView software and saved as TIFF video files. The recording plane covering < 200μm x 200μm at 256×256 pixel resolution was typically ∼50-100 microns deep relative to the upper surface of the slice, to retain good signal strength, whilst ensuring the neurons imaged were less likely to have been damaged during slice cutting. These acquisition parameters, including the decision to image a single plane rather than a volume, were chosen in order to allow adequate cellular spatial resolution whilst covering a field-of-view encompassing most of the LC core. The modest temporal resolution was judged acceptable due the slow temporal dynamics of GCaMP6s with a rise time of ∼200ms, decay time of ∼600ms for a spike evoked transient (Chen et al., 2013). Each recording ended with a high potassium aCSF wash (in mM: NaCl 77, KH2PO4 1.2, KCl 51.9, NaHCO3 26, D-glucose 10, CaCl2 2.4, MgCl2 1.3) to depolarise GCaMP6s positive cells to identify healthy, responsive neurons for subsequent post-hoc analysis.

Initial imaging data pre-processing was performed in FIJI using built in functions (Schindelin *et al*., 2012). Firstly, correction of any X-Y drift of the slice during recordings was performed using a process of image registration. The *’despeckle’* function (a 3×3 pixel median filter) was used to remove ‘salt & pepper’ noise and aid subsequent motion correction. Registration of time series images was performed using *’Template Match’* function to align all images in the time series to a reference image, usually chosen from the half-way time point of the recording. The average fluorescence values over time for individual LC cells was extracted. To do this, first a reference image for cell segmentation was produced using a time-projection of the final high potassium wash segment of the recording (*Z-project function*), in order to visualise the cells exhibiting high fluorescence deltas in response to high potassium wash. Using this image, cell bodies were manually segmented as ‘regions of interest’ (R.O.I.s) using the *R.O.I. manager*, and fluorescence trace of each ROI was extracted as the average pixel intensity at each time point, along with the X-Y location within the image.

Following the generation of these time-series traces, subsequent analysis was performed in Python. In slices where z-plane drift had occurred during recordings (visible as slow, correlated shifts in signal), drift correction was performed, using a 3rd order polynomial fitted and subtracted from the raw fluorescence trace, as this method was able to correct drift whilst preserving dynamic responses e.g. to high potassium wash (Figure S3). To reduce high frequency noise, data was low pass filtered using a Savitzky-Golay (SG) filter (Savitzky & Golay, 1964) – a digital filter designed to remove noise by smoothing data without distorting the underlying signal, by fitting successive sub-sets of adjacent data points with a low order polynomial by a linear least-squares method. The SG filtering was performed with a window size of 25 time points and a 3rd order polynomial, from Scipy Signal toolkit, after initial pilot uses showed these parameters to smooth high-frequency imaging noise, whilst preserving the underlying GCaMP6s responses e.g. to high potassium (Figure S3).

For figures showing initial data processing the units used will be typically be ΔF/F_0_ :

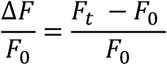

F_0_ = average fluorescence over the initial baseline 200s of the recording

F_t_ = Fluorescence at a given time point.

Defining F_0_ according to the first 200s of recording allows quantification of GCaMP6s transients, providing the basal fluorescence remains stable over the recording. However, these measurements can become less reliable if measuring faster transients, occurring at different time points relative to slower changes in baseline fluorescence. To counter this, F_0_ can be calculated as a shifting baseline, calculated using a rolling window (Romano *et al*., 2017). However, using this method to define F_0_ during ΔF/F_0_ standardisation can introduce artefacts - shown using data from zebrafish cerebellar and hindbrain neurons responding to vestibular stimuli including reductions in fluorescence signal relative to baseline F_0_ (Vanwalleghem *et al*., 2020). Using a moving F_0_ baseline on the data in our study and transforming traces into ΔF/F_0_ also resulted in artificial positive transients following inhibitory responses, some reaching a similar magnitude to expected positive transients in GCaMP6s signal. This distortion was avoided by the use of Z-scoring to transform ΔF time-series data. As such, the majority of GCaMP6s signal changes and time-series plots will utilise Z-scoring to standardise individual neuronal responses in terms of their standard deviations (resulting in changes in fluorescence value reported as σ):

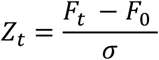

where σ = standard deviation of the fluorescence initial 200 seconds of recording

Following ROI extraction and pre-processing, neurons responsive to high potassium were identified using a method for detecting significant responses in GCaMP6s signal in response to pharmacological stimuli (Figure S4). To generate bounds representing plausible noise around the basal fluorescence signal, a Gaussian model was fitted to the first 200 seconds of the recording (a period without any interventions), and used to generate confidence intervals for each individual neurons’ activity, above and below the pre-intervention baseline (the average value over the 50 seconds preceding the intervention – typically a drug application). Cells were identified as showing significant responses when the GCaMP6s signal remained outside the 90% confidence interval following the intervention, for a pre-specified period depending on the expected length of the response (ie depending on the drug kinetics/dynamics of action), and were classified as excitatory or inhibitory. For example, potassium responsive cells were defined as those in which excitatory transients lasting 30 seconds or longer were detected above a 90% confidence interval in the 200 second window following entry of high potassium aCSF into the perfusion system.

A variant of this method was used to detect significant PSEM308 responses, with the addition of a dynamic baseline around which the confidence intervals were centred. This is based on that described in the Romano toolkit (2017) but adapted due to the specific requirements of GCaMP6s imaging in LC neurons. First, a rolling baseline (fSmooth, Figure S5A) was calculated, aiming to capture slow basal changes in fluorescence, but not the transients evoked by drug application. fSmooth was calculated as the median value of a rolling window (300s, 500x larger than the decay constant of GCaMP6s (0.6 s (Chen et al., 2013)). In pilot analysis, this fSmooth value was found to produce a baseline that adequately captured slower changes in GCaMP6s signal, whilst allowing PSAM-mediated excitatory transients to be detected (Figure S5B). The rolling window size is due to the fact that the responses we are aiming to detect are slower responses (taking place over 10s-100s of seconds), resolved using a relatively low (∼1Hz) imaging frequency. As LC neurons exhibit tonic, pacemaker firing, and GCaMP6s signal under basal conditions reflects that tonic firing (see figure S6). With the fSmooth baseline calculated in this manner, a Gaussian noise model was fitted to the first 200 seconds of recorded activity for each neuron (Figure S5B). This could then be used to classify neurons as showing excitatory responses to PSEM308 if they exceeded the 90% confidence interval around fSmooth for at least 30 seconds in the 40-160 seconds following PSEM308 entry into the perfusion system (Figure S5C), or as showing inhibitory responses if they fell below the 90% confidence interval or non-responsive (Figure S5D). For detection of neurons showing biphasic responses (inhibition following excitation, or vice-versa), significant responses were defined as those which exceeded the confidence bounds for a minimum of 30 seconds in the 200-350 seconds following PSEM308 entry into the perfusion system.

### Computational modelling

#### LC neuron model

The starting point of the computational modelling performed here was a Hogdkin-Huxley formalism, empirically grounded LC neuron model which was first described by Alvarez et al (2002). The model consists of two compartments (soma and dendrite), in which the rate of membrane voltage is described by differential equations which depend on the sum of ionic conductances (all currents are given in µA/cm^2^ of membrane). In addition to the standard fast voltage activated sodium and potassium conductances, the Alvarez model also has a high voltage activated calcium conductance, a calcium activated after-hyperpolarising current and a persistent sodium conductance as well as a leak current. A model of intracellular calcium was also implemented with a dependency on membrane voltage but without formal spatial representation or buffering. This implementation generated spontaneous discharge of action potentials with a relatively pronounced AHP. The original Alvarez model was re-coded in Python3 (GITHUB LINK) and it is described in appendix 1 and only the substantive extensions to the model are outlined below.

### Extensions to Alvarez model and simulations

#### α2R model

Noradrenaline, released following action potentials binds to pre- or post-synaptic α2 receptors which activates G-protein activated inward rectifying potassium channels (GIRKs), allowing outward flow of potassium ions causing membrane hyperpolarisation. To model this process, an alpha-function waveform was chosen as a simple, computationally efficient method, with established use as a general model of synaptic inhibition (Ermentrout & Terman, 2010) as well as specific use as a model of α2R inhibition in LC neuron populations (Patel & Joshi, 2015). The alpha-function waveform was used to describe a transient increase in an inhibitory conductance, *g*_α2_, following each spike, with a time constant chosen with reference to experimental literature describing long (100s of milliseconds) transients for α2R mediated signalling in the LC (Egan *et al*., 1983; Courtney & Ford, 2014). For *n_j_* spikes arriving at neuron *j*, with the *r^th^* spike arriving at time *t_j_ ^r^*, then for neuron *j* the inhibitory conductance *g*_α2_ is given as:

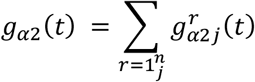

Where:

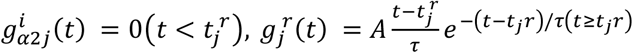

*τ* = time constant, 400ms by default

A = scaling factor = 0.3

*g_α2_* then determines a potassium current as follows:

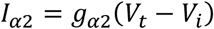

Where: *V_i_ =* -104mV - conductance of α2R mediated *K^+^* conductance in LC (Egan *et al*., 1983)

#### Modelling opto-activation and lateral inhibition

Used simulations of pairs of LC neurons in which opto-activation of one ‘ChR2-transduced’ neuron was modeled via 10ms pulses of high current injection, calibrated to drive one-to-one entrainment action potential discharge. This neuron was connected to a partner via an α2R synapse to assay the inhibitory response in the neighbouring neuron (as described above).

#### Calcium-dependent noradrenaline release

Vesicular release of NA is known to be a calcium dependent process, with intracellular calcium levels governing spike-related release (Courtney & Ford, 2014). As the LC neuron model includes terms to describe intra-cellular calcium (including transient spike-related increases), the basic alpha-function model describing lateral inhibition was augmented for some simulations (detailed in the results text and figure legends) to introduce a calcium dependence to the magnitude of this inhibition.

Vesicular release is regulated by the extensive family of synaptotagmin proteins, hypothesised to act as Ca^2+^ sensors governing exocytosis (Tucker & Chapman, 2002). Intracellular Ca^2+^ increases within hundreds of microseconds of action potential generation via the opening of voltage gated calcium channels (VGCCs), binding to synaptotagmin proteins and activating them, leading to subsequent activation of the membrane fusion machinery including SNARE proteins (Sudhof, 2012). A general kinetic model for calcium dependent vesicular release requiring synchronous binding of release proteins in this manner, has been developed, known as the synchronous release pathway (Nadkarni *et al*., 2010). In this model, the scalar A for α2R release events is dependent on a finite pool of two-state (active or inactive) release proteins, with each release protein requiring sequential binding of multiple Ca^2+^ ions for activation:

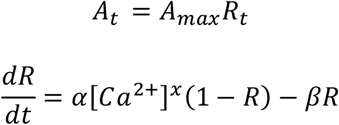

Where:

α = Ca^2+^ association constant

β = Ca^2+^ disassociation constant

R = proportion of release proteins in active state (between 0 -> 1)

*x* = Ca^2+^ binding exponent = 5, representing the number of Ca ions required to simultaneously bind for release proteins to move to active state (as per Nadkarni et al (2010)).

#### Modelling biphasic PSAM responses

Based on the known biological mechanism of PSAM activation, i.e. the opening of direct ligand gated ion channels to drive membrane depolarisation, PSAM activation was approximated using an alpha function to describe a transient increase in an excitatory sodium conductance *g_PSAM_*: 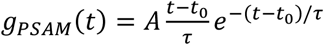

where:

*t_0_* = time of PSAM onset

*τ* = time constant = 2000msec

*A =* amplitude scalar = 0.2

This allows the PSAM mediated current to be determined by:

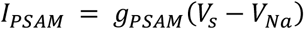

leading to the following equation for membrane voltage in PSAM-transduced neurons:

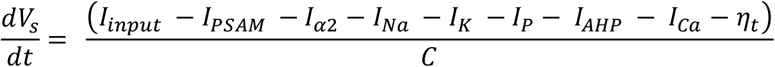

For simulations investigating biphasic α2R mediated responses, the alpha-function describing *g*_α2_ transients was replaced with a double exponential, to allow specific alterations to be made to the decay time constant without affecting the rising phase of the transient. For *n_j_* spikes arriving at neuron *j*, with the *r^th^* spike arriving at time *t^r^_j_, g*_α2_ is given as:

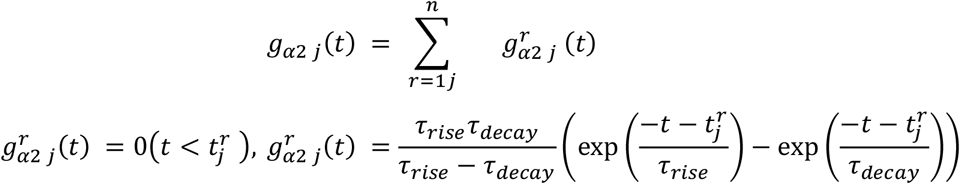

Where:

*τ*_*rise*_ = rise constant = 100msec

*τ*_*decay*_ = decay constant

To model the effect of NET reuptake saturation on α2R transients, a sigmoid transfer function was used to derive *τ*_*decay*_using *g_α2_* as a proxy for the level of noradrenergic tone at the time of each release event. The sigmoid Hill-Langmuir equation describes the binding of ligands to macromolecules, and in this case described how at low levels of released NA, efficient reuptake by NET from the extracellular space into LC neurons would promote the discrete temporal fidelity of synaptic events. However, as the concentration of released NA rose, NET molecules begin to become saturated, resulting in less efficient NA reuptake and a prolongation in the time-course of synaptic events, eventually peaking as maximal levels of NA release per spike were reached.

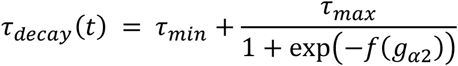

Where

*τ*_*min*_= lower bound for *τ*_*decay*_ = 200msec*

*τ*_*max*_ = upper bound for *τ*_*decay*_ = 4000msec*

*f*(*g*_*α*2_) = *g*_*α*2_ transformed by linear interpolation as follows:

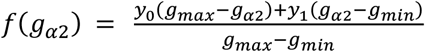 where:

*y_0_ =* minimum input value for sigmoid function = −10

*y_1_ =* maximum input value for sigmoid function = 10

*g_min_ =* peak *g_α2_* observed during basal firing = 2e^-08^

*g_max_ =* peak *g_α2_* observed during maximal (50Hz) firing = 0.1

* - chosen with reference to Courtney & Ford (2014)

### Histology

Animals were terminally anaesthetised with pentobarbitone (200mg/kg, i.p), and trans-cardially perfused with 300ml 0.1M phosphate buffer solution (PBS), followed by 300ml 4% formalin. Brains were removed and post-fixed in 4% formalin overnight before being placed in 30% sucrose solution until saturated. Slices were cut using a freezing microtome (30μm). For immunohistochemistry, slices were incubated with primary antibodies at 1:2,000 dilution for 12 hours, then with secondary antibodies at 1:500 dilution for 3 hours, with 3 sequential washes in 0.1M PBS after each antibody (see Table 3). Slides were mounted in Fluorosave medium on glass slides, and imaged on a fluorescent microscope, (Leica DM4000B using LAX suite software). Cell counting was performed manually using FIJI’s *Cell Counter* plugin (Schindelin *et al*., 2012), with cells defined as those with a visible nucleus and appropriate size and morphology for LC. Vector efficacy was defined as the percentage of noradrenergic LC cells (labelled using Dopamine beta-hydroxylase (DBH) antibodies) that also expressed the virus as marked by a fluorophore. Vector specificity was defined as the proportion of transduced cells that were also noradrenergic LC cells.

**Table 3.** Antibodies used for histology.

| ***Primaries*** |  |  |  |
| --- | --- | --- | --- |
| ***Target*** | ***Host*** | ***Concentration*** | ***Supplier & ID*** |
| *DBH* | *Mouse* | *1:4000* | *Millipore MAB308* |
| *GFP (GCaMP6s)* | *Rabbit* | *1:4000* | *Invitrogen A11122* |
| ***Secondaries*** |  |  |  |
| ***Fluorophore*** | ***Host & Target*** | ***Concentration*** | ***Supplier & ID*** |
| *AlexaFluor594* | *Goat anti-mouse* | *1:1500* | *Life-Tech A11032* |
| *AlexaFluor488* | *Goat anti-rabbit* | *1:1000* | *Life-Tech A11008* |

## Results

Whole-cell recordings were made from LC neurons in pontine slices cut from rats that had direct stereotaxic injection of CAV-PRS-ChR2-mCherry. This enabled optogenetic stimulation by an LED fibre placed above the LC (Figure 1A, as previously described in Li et al., (2016)). In transduced LC neurons, this optoexcitation caused the expected large inward currents in voltage clamp recordings (Figure 1B). In current clamp, brief pulses of light (10-30ms) were able to entrain action potential firing (Figure S1A). The light stimulus response function for transduced neurones showed a reliable ability to induce firing at physiological rates, with a degree of accommodation seen at frequencies >20Hz (Figure S1). Recordings from non-transduced LC neurons (in the same slices but without mCherry fluorescence) never showed light-evoked action potential discharge (n=10).

**Figure 1.**
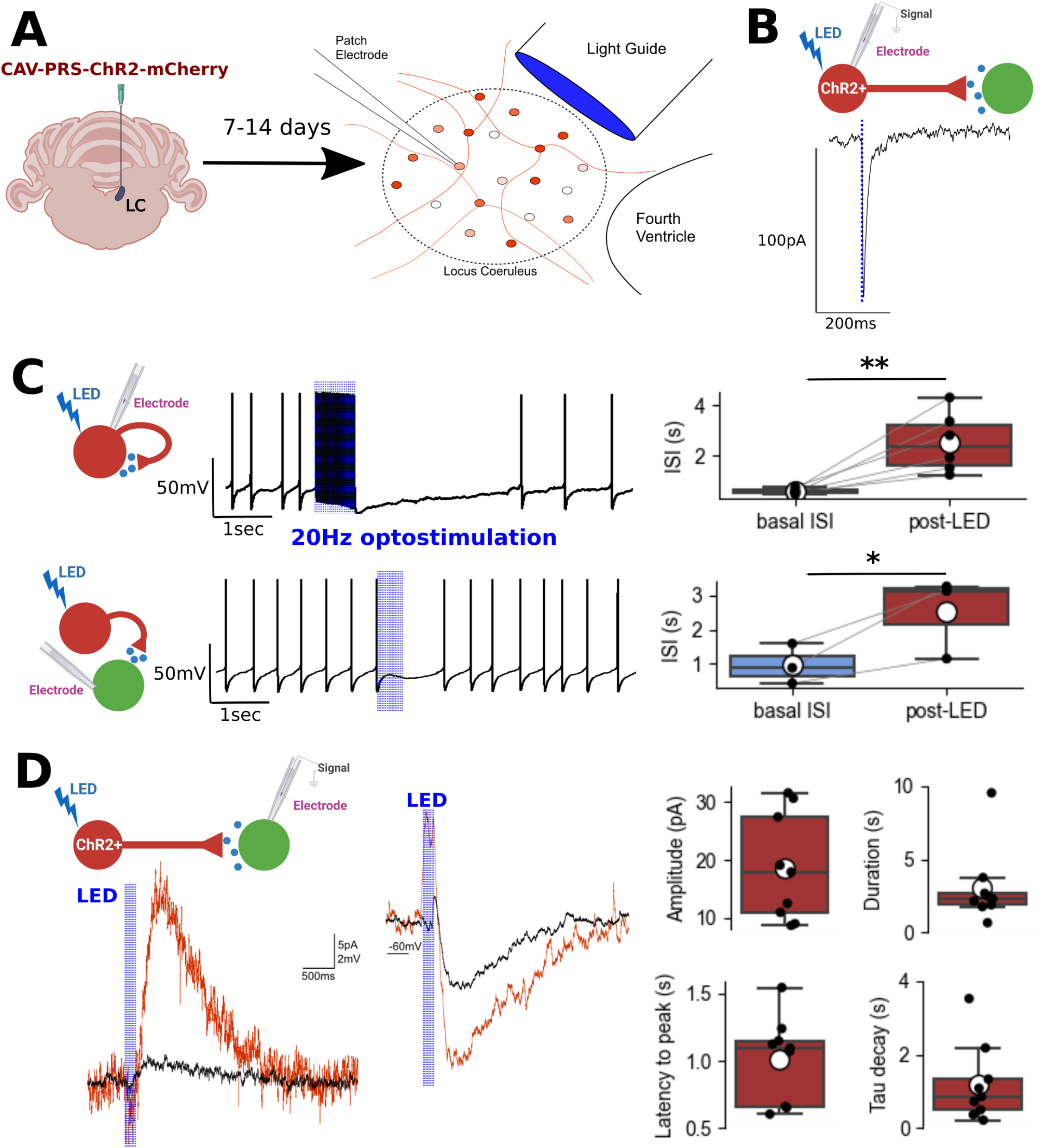
Optoactivation of LC neurons, and evoked inhibition. A – Ex-vivo LC slices were generated following direct targeting of the LC nucleus with CAV-PRS-ChR2-mCherry. Whole-cell recordings were made from LC neurons (both ChR2-transduced and non-transduced) with 470nm light stimulation. B – In ChR2 transduced cells, blue light pulses (30msec) evoked fast inward currents. C – In ChR2-transduced neurons, opto-evoked action potential discharge (Blue vertical lines, 20Hzx30ms) was followed by a period of inhibition of firing (Top panel) that was seen in all recorded cells (plot on right shows change in inter-spike interval for n=6 neurons, 2 tailed paired t-test, ** - p<0.005). A similar phenomenon was seen in non-transduced LC neurons, light flashes (Blue vertical lines, 20Hzx30ms) caused a transient suppression of ongoing spontaneous firing (lower panel), plot on right shows change in inter-spike interval, n=3 neurons, paired 1-tail t-test, * - p<0.05). D – Optogenetic stimulation of voltage clamped neurons also produced slow inhibitory outward currents (left panel) that correspond to the transient membrane hyperpolarisation seen in current clamp recordings in the same cell (middle panel). Responses to single light pulses (average of ten trials) shown in black, and individual responses to a 10×50Hz pulse train shown in red. Right panel plots show summary statistics of the properties of these slow outward currents (n=9 neurons). Box plots – median and interquartile range, whiskers – 1.5 x IQR, white circle – mean.

### Optoactivation of the LC causes an α2-adrenoceptor mediated inhibition

This experimental slice preparation allowed the analysis of inhibitory responses seen to follow optoactivation of LC neurons, both in ChR2-mCherry transduced neurons, and in neighbouring non-transduced LC neurons. A standard optoactivation protocol of a train of light pulses (30ms) at 20Hz for 1 second was employed. In current clamp recordings of transduced neurons, opto-stimulation of the LC produced an inhibition of spontaneous firing following the initial train of evoked spikes (Figure 1C, the inter-spike interval increased from 0.58±0.11 to 2.52±1.19 seconds, n=6, p = 0.0043). In a sparser set of recordings from non-transduced neurons, there was also evidence of an inhibition of spontaneous cell firing after the same illumination protocol but in the absence of any preceding opto-evoked spikes (Figure 1C, the inter-spike interval increased from 0.96±0.59s to 2.5±1.2s, an increase of 185%, n=3, p = 0.042). This is consistent with the concept of collateral inhibition within the LC (Aghajanian *et al*., 1977; Ennis & Aston-Jones, 1986), originally identified using extracellular recordings combined with antidromic activation in rats *in vivo*. A corresponding slowly developing, prolonged outward current was observed in these non-transduced neurons following optoactivation of the LC (Figure 1D, n=3). A similar current was also evident in the ChR2-transduced LC neurons (n=6) which were now prevented from firing opto-evoked action potentials by voltage clamp at −50mV. This finding of slow inhibition without preceding action potential discharge in the recorded neuron is also consistent with collateral inhibition from opto-evoked action potential discharge in neighbouring ChR2-transduced LC neurons. The underlying inhibitory (outward) current had an average amplitude of 18.7±9.2 pA and peaked after 1.01±0.32s with a decay time constant of 1.20±1.06s (Figure 1D). The inhibition could be repeatedly evoked by the 20Hz optoactivation without sign of desensitisation or run down (−11.7±0.6pA initially vs −11.6+0.6pA after 20 minutes, n=5, Figure S2A). The inhibition could also be evoked by a single light flash (Figure 1D) consistent with it being a synaptic event. No slow excitatory responses that persisted after the immediate period of optogenetic stimulation were seen in any LC neuron (ChR2-transduced or non-transduced).

To test whether these inhibitory responses were mediated by the release of NA and subsequent activation of α2R, we examined the effect of a selective α2R antagonist (atipamezole, 5μM) and a selective NA reuptake inhibitor (reboxetine, 10μM) on the evoked outward currents in recordings from transduced neurons. The outward current was blocked by atipamezole (from −12.6±1.6pA in controls to −3.8±0.4pA, P<0.0001, n=6, Figure 2A). Application of atipamezole also produced a shift in the holding current (11.0±6.6pA compared to baseline, n=6) consistent with the blockade of a tonic α2R-mediated inhibitory current produced by ongoing spontaneous NA release within the LC. The slow outward current was reversibly enhanced by reboxetine (n=7, Figure 2B) in both amplitude (−9.0±1.2 to −14.7±1.3 pA, P<0.001) and duration (5.7±0.9 to 22.4±2.7 seconds, P<0.001).

**Figure 2.**
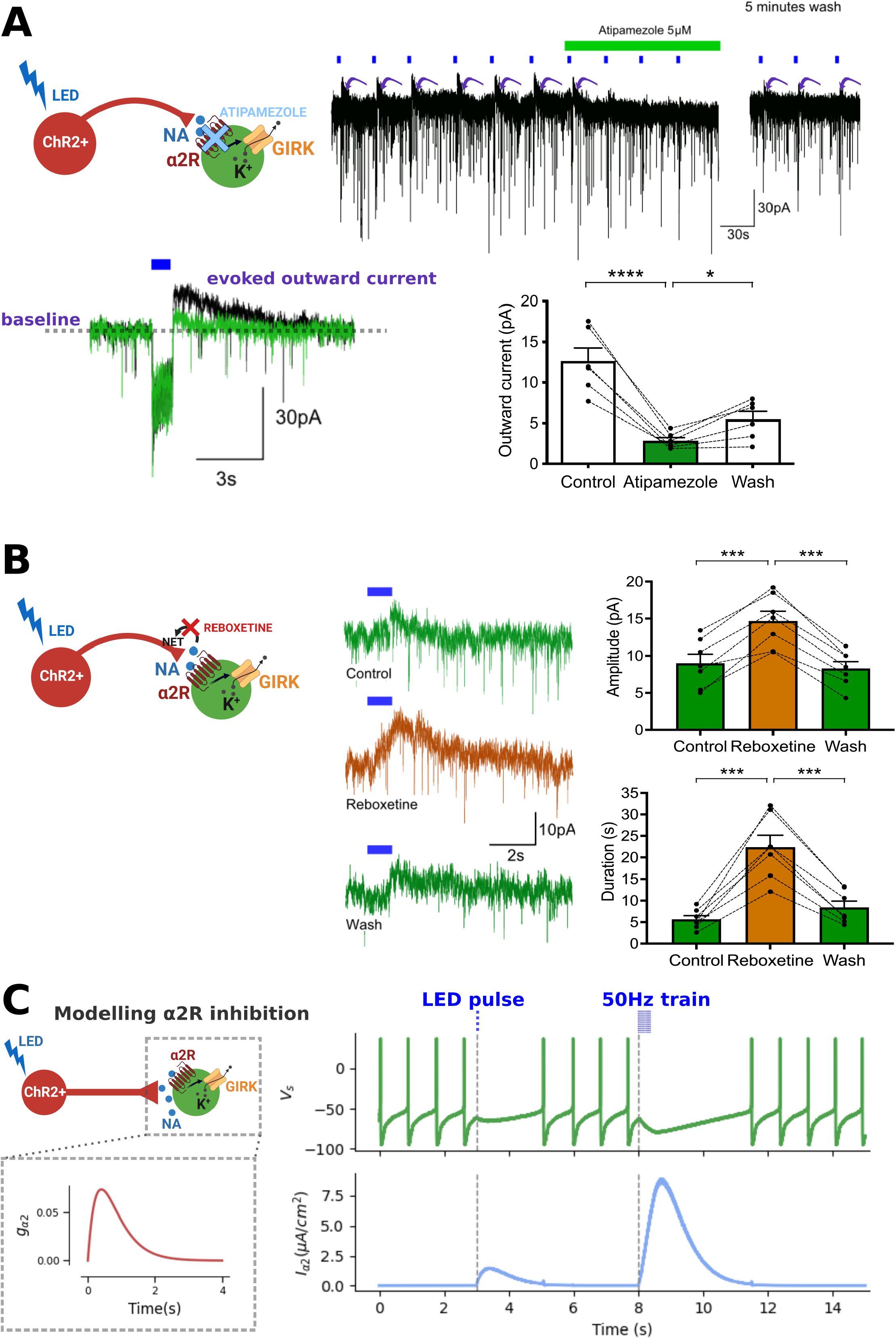
Slow inhibition is noradrenergic and mediated via α2Rs. A – Outward currents following opto-stimulation were reversibly blocked by the α2R antagonist atipamezole. 20Hz optostimulation was administered every 30 seconds, followed by atipamezole (5µM, top panel, bottom left panel – black trace shows baseline outward current and green trace shows current in the presence of atipamezole). Atipamezole attenuated the outward current which partly reversed on washing. (Bottom right, RM ANOVA, F = 42.85, DF=2, n=6, p<0.0001, with Holm-Sidak’s post-hoc testing, **** - p ≤ 0.0001, * - p <0.05) B –The noradrenaline reuptake inhibitor reboxetine (10µM) increased the amplitude and duration of outward currents (middle panel shows example recordings). Reboxetine reversibly increased the amplitude of outward currents (top right, RM ANOVA, F = 53.96, DF=2, n=7, p<0.0001, with Holm-Sidak’s post-hoc: *** p < 0.001) as well as reversibly increasing duration (bottom right, RM ANOVA, F = 52.41, DF=2, n=7, p = 0.0002, with Holm-Sidak’s post-hoc *** - p<0.001) C - Computational modelling of NA release following optoactivation, using an alpha-function to describe transient GIRK channel activation (inset left panel), was able to replicate the inhibitory effects of opto-stimulation on neighbouring (ChR2 negative) neurons on basal firing.

To investigate the ionic basic of the evoked inhibitory current, we made recordings using a pipette solution containing CsSO_4_ (100mM) and Tethraethylammonium (5mM) to block K^+^ channels and made a comparison with recordings using the standard K-Gluconate-based solution (Fig S2A). After obtaining the whole-cell recording configuration, the typical profile of slow inhibitory currents were observed following optoactivation (10.8±1.6pA for Cs-TEA, n=5, Figure S2). However, in the presence of Cs-TEA internal solution this current progressively reduced in amplitude over a period of 15 minutes (to 1.5±0.3pA vs 11.6±1.2pA K-Gluconate, P<0.0001). This slow noradrenergic inhibitory conductance also reversed at a membrane potential of −110mV and showed inward rectification (Fig S2B) consistent with previous reports of α2R-mediated activation of potassium conductance in LC neurons (Williams *et al*., 1985).

These experiments indicate the slow current was caused by optostimulation-evoked noradrenaline release producing an inhibitory post synaptic current (α2R-IPSC) and that there is an ongoing background release of noradrenaline in the LC in the absence of optoactivation that likely reflects spontaneous tonic firing. This process was modeled in paired-neuron simulations, with one neuron ‘optogenetically’ driven to fire at specific frequencies producing α2R-IPSCs in the paired postsynaptic neuron generated using an alpha-function to model transient increases in the inhibitory conductance *gα2* (Figure 2C). This was sufficient to qualitatively mimic the opto-evoked lateral inhibition of spontaneous firing, as observed in experimental data (Figure 1C-D).

### Properties of α2R-inhibitory postsynaptic currents in the LC

Additional optogenetic experiments were undertaken to investigate the relationship between presynaptic firing and the α2R-IPSCs. The dependence of opto-evoked NA release on propagating sodium spikes was investigated using bath applied TTX (500nM). This did not fully block spike discharge in LC neurons (Figure S3A, n=5). The remaining spontaneous spikes were smaller (reduced from 81.1±2.3 to 43.3±2.1mV) and had increased duration (1.2±0.1 to 4.5±0.7ms) consistent with them being calcium spikes as previously reported (Williams *et al*., 1984; Williams & Marshall, 1987). TTX attenuated the α2R-IPSC by 25% (Figure S3B-C, from −31.1±6.3 to −23.3±5.1pA, n=5, p=0.013), but did not abolish it, indicating that NA release and the subsequent inhibition can occur in the absence of sodium-based action potentials and suggesting that it may also be somato-dendritic rather than solely axonal in origin (Huang *et al*., 2007). We therefore termed this lateral inhibition that may be due to either somato-dendritic release or feom LC axon collateral terminals.

To explore the relationship between stimulation frequency and the α2R-IPSC we measured the charge transfer in response to a fixed number of light pulses (10 pulses x 10ms) at varying frequencies (10-50Hz, Figure 3). Either side of the standard 20Hz stimulus-evoked outward current (shown in Figures 1&2) there was an exponential change in the amplitude of the α2R-IPSC, which was well fitted by a single exponential (Figure 3B, R^2^ = 0.92). This steep relationship between IPSC amplitude and increasing firing frequency was not a consequence of an increase in the absolute number of presynaptic action potentials evoked by the optostimulation as direct recordings from ChR2-transduced LC neurons showed that the number of spikes tended to reduce slightly with a degree of accommodation at stimulation frequencies >20Hz (see Figure S1B). This suggests the frequency dependence of the α2R-IPSC is either due to changes in NA release, reuptake or receptor activation at different stimulation frequencies.

**Figure 3.**
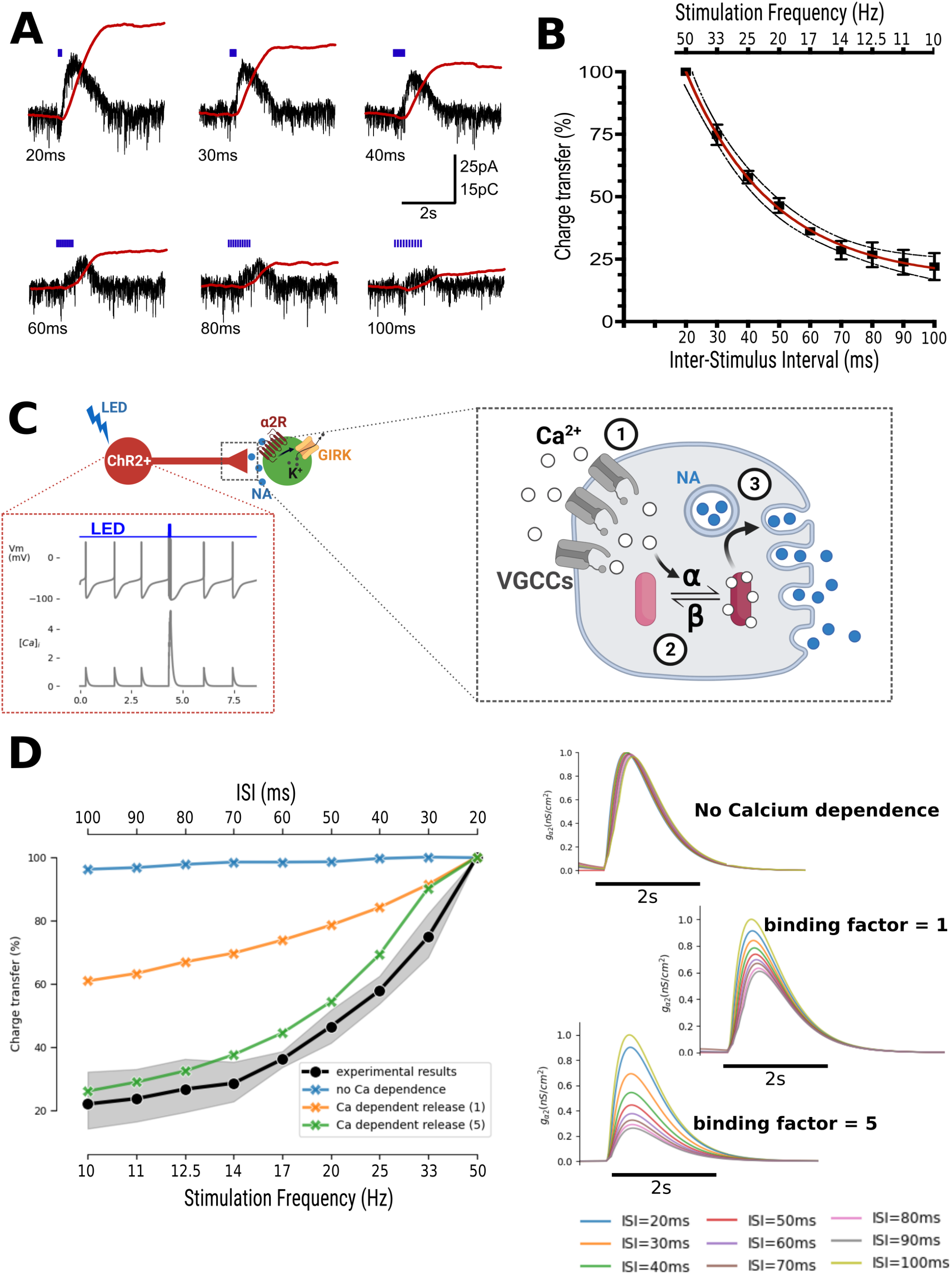
α2R-IPSCs show strong non-linear frequency dependent facilitation. A – α2R-IPSCs show a strong dependence on opto-stimulation frequency (example responses shown for inter-stimulus intervals of 20-100ms. Red traces show charge transfer calculated as area under the curve). B – The relationship between inter-stimulus interval and charge transfer was well fitted by a one-phase exponential (mean ± SEM, n=5, dotted lines - 95% confidence interval of curve, R^2^=0.92) C - Paired neuron simulations, in which one neuron was driven to fire at specified frequencies, replicating optoactivation (inset red box), coupled to a synchronous pathway model of calcium dependent NA release (detailed in inset grey box, right panel). Step 1 action potentials open voltage gated calcium channels (VGCCs) leading to intracellular calcium rises (summation of calcium transients during high frequency stimulation shown in inset trace - red box). Step 2 Intracellular calcium binds to and activates release proteins, with simultaneous binding of up to 5 calcium ions required to activate each release protein. Step 3 Activated release proteins facilitate exocytosis of vesicular NA. D - This synchronous pathway mechanism replicates the non-linear relationship between optogenetic stimulation, especially when the number of calcium ions required for activation of release proteins increases above 2, in way that paired neuron models without calcium-dependent release fail to do. (ISI = inter-stimulus interval). Left panel shows the relationship between stimulation frequency and charge transfer for paired neuron simulations with/without calcium-dependent release, with calcium-dependent release showing similar frequency dependent inhibition to experimental results (experimental data mean +/- 95% confidence interval, n=5). Right panel shows transient increases in the inhibitory conductance gα2 generated at different ISIs, for modelled pairs with no calcium dependence release, and for calcium binding factors of 1 and 3 (traces are normalised relative to the peak of the response to 50Hz stimulation).

### Calcium-dependent NA release model replicates the non-linear frequency dependence of α2R-IPSC

The potential mechanism underlying the frequency dependence of the α2R-IPSC was explored using modelling. The initial simple simulation of the synaptic event governed by an alpha function (Figure 2C) did not show the steep frequency dependent facilitation (Figure 2C and 3D). It has previously been shown that NA release from LC neurons occurs in a calcium-dependent manner (Egan *et al*., 1983; Huang *et al*., 2007; Courtney & Ford, 2014; Feng *et al*., 2019). Simulations of optogenetically-evoked lateral inhibition were performed with calcium-dependent release model ((Nadkarni *et al*., 2010) and see methods). The best approximation to the observed data was provided by a model with 5 calcium ions binding co-operatively to the release machinery which closely replicated the non-linear dependence of the α2R-IPSC on presynaptic firing frequency (Figure 3C&D). This supports the hypothesis that lateral inhibition in the LC can be understood as a somato-dendritic phenomenon relying on activation of vesicular release proteins by calcium.

### Developing a LC network recording strategy with genetically-encoded calcium indicators

The optogenetic patch-clamp recording experiments demonstrated the existence of inhibitory, α2R-mediated interactions between LC neurons. However, a change in experimental strategy was needed to investigate whether the observed α2R-inhibition reflected a non-specific spatial ‘surround’ inhibition or a more structured network interactions such as targeted lateral inhibition. Imaging with a genetically-encoded calcium indicator (GECI) deployed as a methodology, to allow simultaneous recordings of activity from ensembles of LC neurons. A novel canine adenoviral vector construct (CAV-PRS-GCaMP6s) was designed to selectively express a GECI in the LC. Stereotaxic injections targeting the LC using this vector (Figure 4A) achieved a similar level of performance to equivalent CAV-PRS driven vectors (Li *et al*., 2016; Hirschberg *et al*., 2017; Hayat *et al*., 2020), with a transduction efficacy of 64.3±2.5% and a specificity of 86.3±10.1. LC neurons expressing GCaMP6s were visible with two-photon imaging under basal conditions in ex-vivo pontine slices. These fluorescent neurons showed the expected large increases in signal when depolarised by perfusion of high potassium aCSF (50mM, Figure S4). The presence of this response to high potassium aCSF, applied at the end of each recording session, was used as a criterion to identify viable, GCaMP6s-expressing LC neurons in all subsequent recordings.

**Figure 4.**
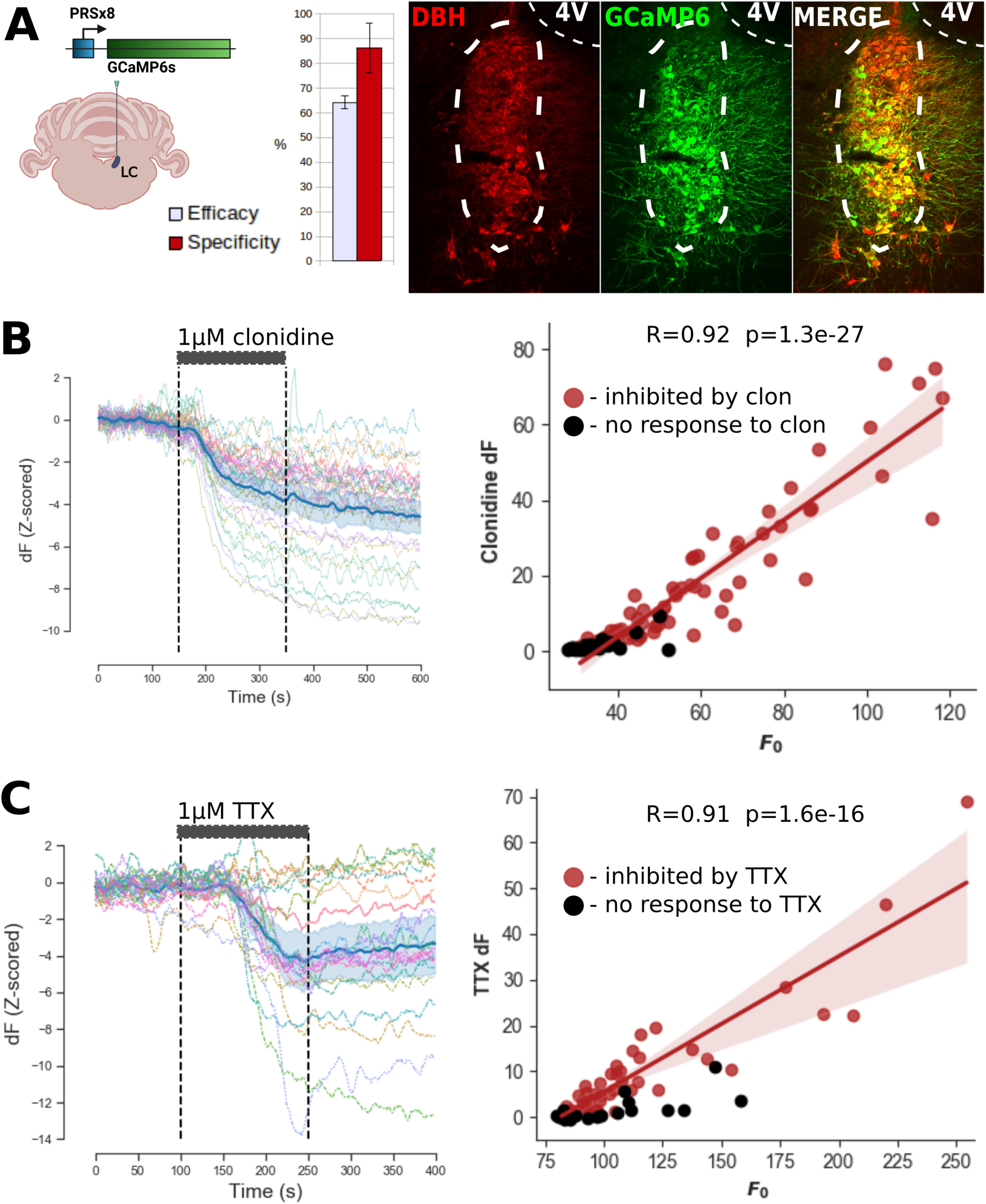
Expression of GCaMP6s in LC neurons allows inhibition to be resolved. A – CAV-PRS-GCaMP6s was injected directly targeting the LC. This resulted in a transduction efficacy of 64.3±2.5% of dopamine-beta-hydroxylase (DBH) positive LC neurons and a specificity of 86.3±10.1% (mean±SD, n=5 sections from 1 rat). Right panels show a representative section, with GCaMP6s expression in DBH positive neurons within the boundaries of the LC (white dashed line). 4V = 4^th^ ventricle. B - Clonidine (1μM) produced a decrease in GCaMP6s signal in a majority of LC neurons. Left panel shows recording from a single slice with multicoloured traces showing individual neuronal responses (n=36 cells, blue line mean ± 95% confidence interval shaded). The magnitude of the clonidine inhibition for each neuron correlated with basal fluorescence (F_0_, n=96 neurons, 4 slices from 2 rats). Line and shaded region show regression line and 95% CI, fitted to clonidine responsive cells (n=67). C - TTX (1μM) resulted in a decrease in GCaMP6s signal in the majority of LC neurons (42/63, 6 slices from 3 rats). Left panel shows data for a single recording with multicoloured traces showing individual neural responses (n=21 cells, blue line indicates mean with 95% confidence interval shaded). The magnitude of the TTX inhibition for each neuron correlated with the basal fluorescence (F_0_, n=42). Linear regression and 95% CI shaded for model, fitted to TTX responsive cells.

### Resolving bidirectional changes in LC activity

Almost all previous studies using GECIs in neurons have described increases in calcium signal associated with action potential discharge with sparse basal firing rates (Vanwalleghem *et al*., 2020). However, a few studies have recently reported being able to resolve inhibition using GECIs in populations of tonically active neurons (Favre-Bulle *et al*., 2018; Gong *et al*., 2020). To determine if inhibition of LC neurons could be resolved using GCaMP6s imaging, a series of recordings was undertaken using pharmacological strategies known to inhibit LC neurons.

Because we were interested in an α2R-mediated inhibitory mechanism in the LC then we explored the effect of the α2R-agonist, clonidine (1μM). Perfusion of clonidine reduced the mean GCaMP6s signal in the majority of LC neurons (n=67/96, 70%, Figure 4B and see Figure S5 for details of significant transient detection). For these clonidine-inhibited neurons, the average magnitude of inhibition was 7.4±4.8 σ. The magnitude of each neuron’s response to clonidine correlated with its baseline fluorescence F_0_ (Figure 4B, Pearson’s R=0.92, p=1.3e^-27^, n=67 neurons). The relationship between the size of the clonidine inhibition to basal fluorescence in each cell is likely a result of differences in basal firing rate of LC neurons, which determines the basal GCaMP6s signal, and therefore the available range for visible inhibition (a cell with a low firing frequency will not show much sign of inhibition because of a floor effect).

Given that the α2R-agonism may be expected to reduce LC action potential discharge and may also potentially independently alter intracellular calcium levels via second messenger signalling then we also sought to see whether directly reducing action potential discharge would be detectable using the GCaMP6s signal. Application of the fast sodium channel blocker TTX (1μM) produced a significant fall in basal GCaMP6s signal in 42/63 neurons (67%). For inhibited neurons, the average magnitude of inhibition was 4.4±2.8 σ. It is known that the effect of TTX on LC cell firing is not uniform, with some LC neurons continuing to spontaneously fire calcium-based action potentials following TTX blockade of sodium-dependent action potentials ((Williams & North, 1985) and see Figure S3). Therefore, the observation that most but not all LC cells showed a fall in intracellular calcium with TTX is likely because of the presence of this population capable of firing calcium spikes. The magnitude of TTX inhibition in individual cells was tightly correlated with the baseline fluorescence of that cell, F_0_ (Figure 4C, Pearson’s R=0.91, P=1.01e-16, measured across all inhibited neurons). As with clonidine, this indicates that observed TTX responses are dependent on basal calcium levels, likely due to differential baseline firing rates between LC neurons. Taken together the TTX and clonidine experimental findings indicate that GCaMP6s fluorescence changes can be used as an indicator of the inhibition of action potential discharge in LC neurons with spontaneous activity.

### Evidence of lateral inhibition in network recordings

In order to activate a population of LC neurons during calcium imaging recordings, an excitatory chemogenetic ionophore (PSAM^L141F,Y115F:5HT3HC^, (Magnus *et al*., 2011)) was expressed using CAV-PRS-PSAM (Hirschberg *et al*., 2017). Direct transduction of the LC with both the GCaMP6s and PSAM CAV vectors allowed dose-dependent excitation using bolus perfusion of the selective PSAM ligand PSEM308 (3-30μM, Figure 5 & S6). Strong excitatory responses to 30µM PSEM308 were observed in a subset of neurons (4.56±3.99 σ, seen in 24/112 neurons (21.4%)). The responses were temporally discrete with a rapid onset as the agonist reached the slice chamber and typically returning to baseline levels of fluorescence within 150 seconds, allowing repeated application of drug within each recording session. Lower doses of PSEM308 produced smaller responses and reduced the proportion of responsive cells, although excitatory responses were still evident with concentrations as low as 3μM PSEM308 (Figure 5D). In control slices, transduced with only CAV-PRS-GCaMP6s, 10μM PSEM308 elicited no significant excitatory responses (n=29 LC neurons, Figure S7). Similarly, the observation that there were non-responsive neurons within slices injected with both vectors (49/112 cells (43.8%), Figure 5D), indicated that the PSEM308 responses reflect chemogenetic excitation via the PSAM receptor rather than off-target effects of the PSEM308 ligand on neurons without the chemogenetic actuator.

**Figure 5.**
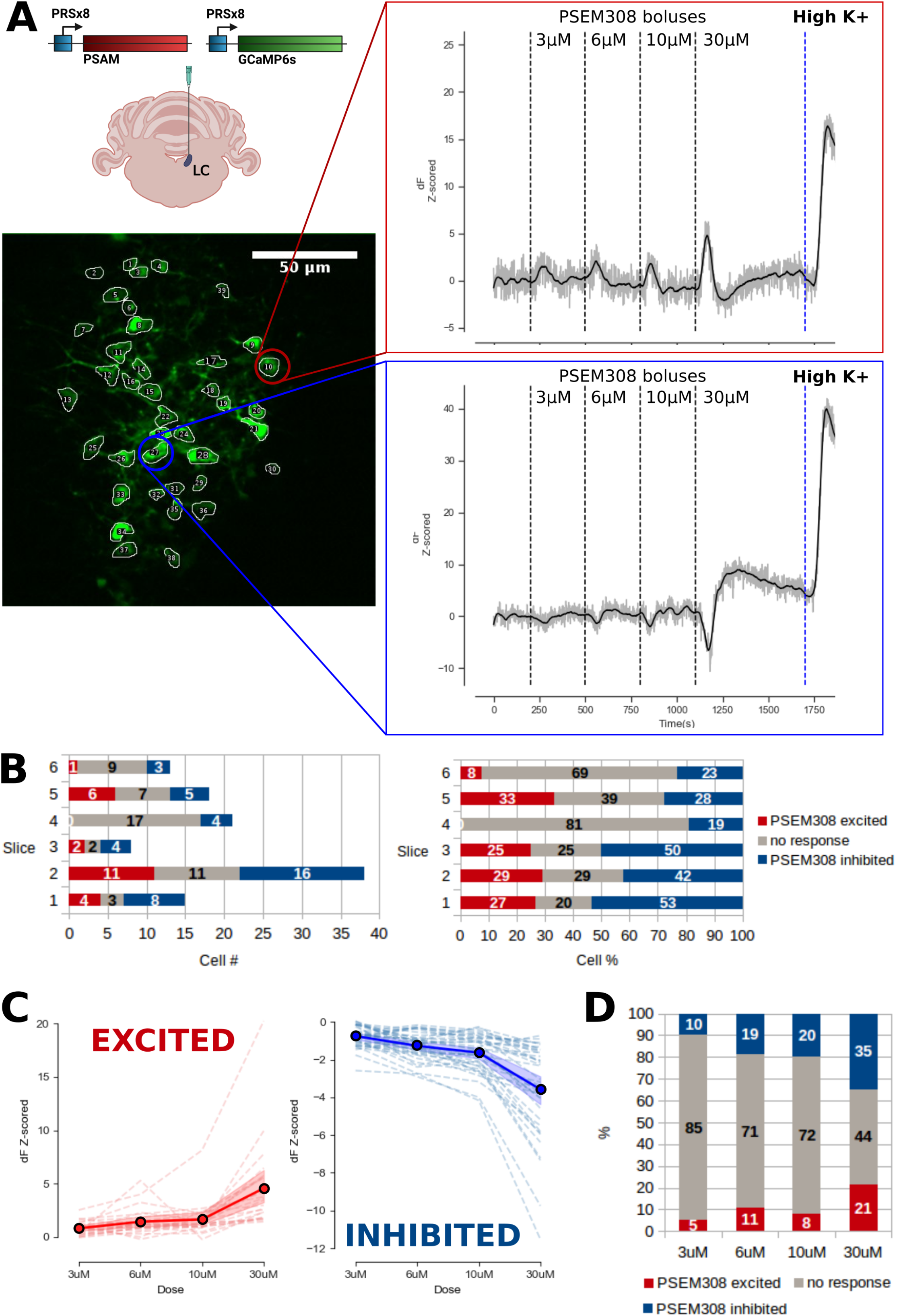
Chemogenetic activation and evoked inhibition of LC neurons. A - Direct injection of CAV-PRS-GCaMP6s and CAV-PRS-PSAM-eGFP targeting the LC resulted in slices with both excitatory (red box) and inhibitory (blue box) responses to increasing doses of the PSAM-specific ligand PSEM308. (Gray trace shows drift corrected data, black trace shows Savitsky-Golay filtered trace used for subsequent analysis). B – Excited/inhibited/non-responsive neurons per slice showing cell numbers (left) and percentage (right) in each response category (6 slices from 3 rats). C – Dose-response for PSEM308 excitation (left panel, n=24/112 cells) and evoked inhibition (right panel, n=49/112 cells). Dashed lines show responses for individual neurons, thick line shows mean ± 95% CI shaded. D – Lower doses of PSEM308 evoked fewer significant responses, but both excitatory and inhibitory responses persisted at 3uM. Plot shows percentage of neurons in each category at different PSEM308 doses for the whole dataset, n=112 neurons.

Within these experiments there was a population of LC neurons which showed marked decreases in intracellular calcium when the chemogenetic agonist was applied (−3.58±2.46 σ, 39/112 cells (34.8%)). These inhibited LC neurons were seen in the same set of slices as the 21.5% (24/112) of neurons which were strongly activated by PSEM308. The excitation and the inhibition had similar timecourses. The magnitude of the inhibitory response depended on the dose of PSEM308 (Figure 5C), and inhibitory responses were present at all PSEM308 doses used (3-30µM), becoming more common as agonist dose was increased (Figure 5D). This corroborated and extended the findings of Hirschberg and colleagues (2017) in a previous study employing the CAV-PRS-PSAM vector using extracellular recording *in vivo* showing both the expected excitatory (50%) and the apparently contradictory inhibitory (25%) firing responses in subsets of LC neurons following local PSEM308 injection *in vivo*.

### Population lateral inhibition is α2R-mediated

As the lateral inhibition was hypothesised to be mediated by α2R-signalling, and to reflect the same phenomenon observed in the single cell optogenetic experiments detailed above, the effect of the α2R-antagonist atipamezole was assessed on the population inhibitory responses. Slices were generated with LC expression of GCaMP6s and PSAM allowing recordings of responses to chemogenetic activation with repeated applications of PSEM308 (10μM), before, in the presence of atipamezole (2μM) and 20 minutes following application (Figure 6A). In those neurons inhibited by the first application of PSEM308 (n=41), a significant effect of drug was seen across the PSEM308 applications (RM-ANOVA, p<0.00005). Atipamezole significantly reduced the inhibitory effect of PSEM308 (Figure 6B, from −1.89±1.08 σ to −1.24±0.77 σ, a reduction of 34.4%, Tukey HSD p=0.0037) and this persisted after 20 mins washout. In contrast, in the same experiments, atipamezole did not block the excitatory PSAM response (Figure 6C, RM-ANOVA, p = 0.0542, n=22 neurons). There was no evidence of diminished excitatory responses with repeated PSEM308 applications for excitation over the course of the recordings. These findings indicate that the lateral inhibition is α2R-mediated and is equivalent to the lateral inhibition seen in the single cell recordings.

**Figure 6.**
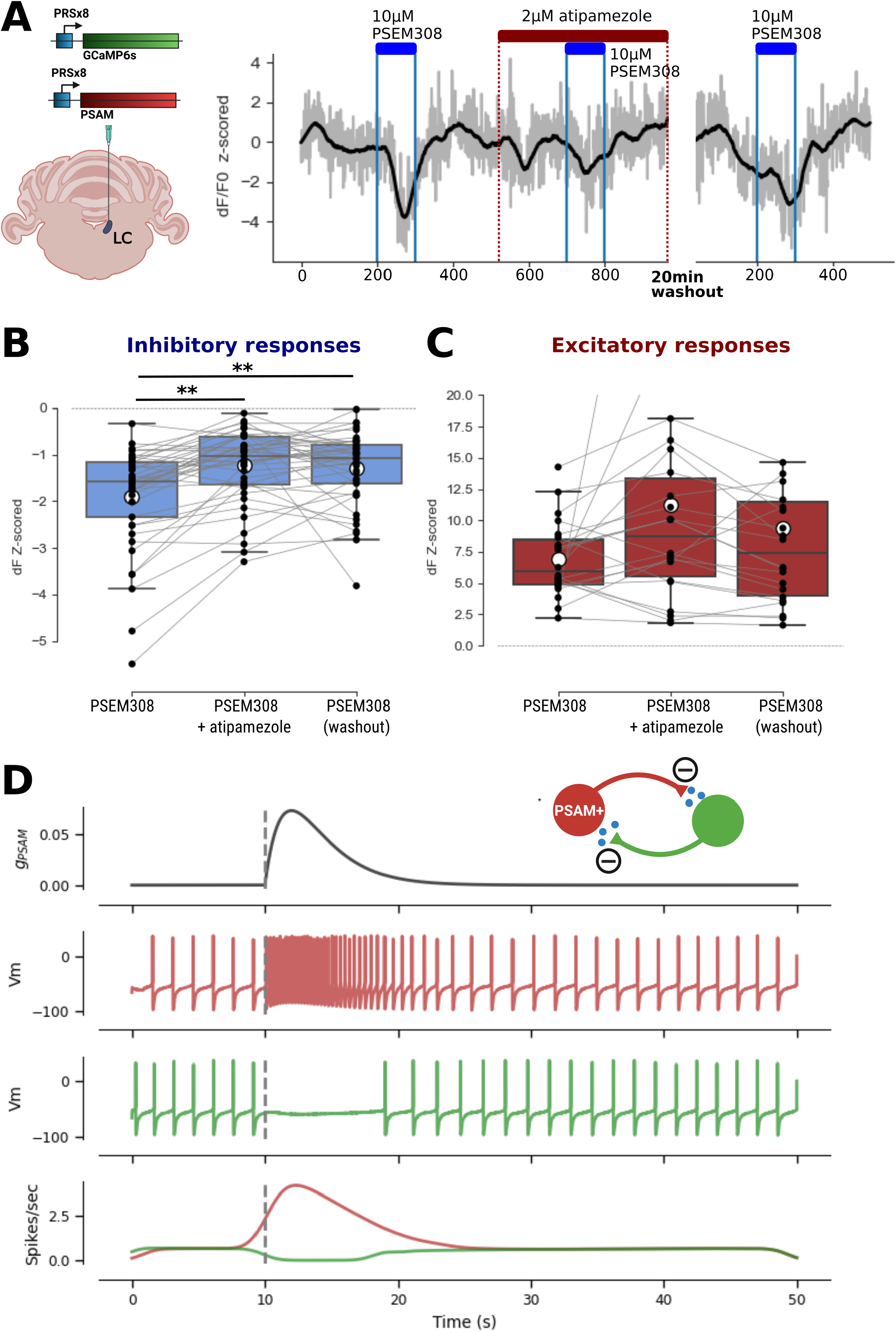
Lateral inhibition is mediated by α2Rs. A – Slice 2P recordings were made following direct transduction of LC with CAV-PRS-GCaMP6s and CAV-PRS-PSAM. Slices were perfused with boluses of PSEM308 (10μM) with/without atipamezole (2μM), then repeated after a 20-minute washout period. Right trace shows the inhibitory responses of a neuron to PSEM308 which was reversibly attenuated in the presence of atipamezole. B –Atipamezole (2μM) significantly reduced inhibitory responses (n=41/151 cells, 27.2%) to 10μM PSEM308. (Boxes show median ± IQR, whiskers 1.5x IQR, mean indicated by white circles, RM-ANOVA, F stat = 15.11, df=2, p < 0.00005, Tukey HSD, ** - P<0.01) C - PSEM308 excitation was not attenuated by atipamezole (n=22/151 cells, 14.6%, RM-ANOVA, F stat = 3.13, df=2, p = 0.0542). (Boxes show median ± IQR, whiskers 1.5x IQR, mean indicated by white circles, Two outlier responses have been cut off to better illustrate the data). D – Paired neuron simulation showing transient activation of a ‘PSAM transduced neuron’ (top trace shows time course of PSAM conductance) was sufficient to generate lateral inhibition in the non-transduced neuron, via reciprocal noradrenaline release and activation of α2Rs.

### Lateral inhibition is insensitive to blockade of amino acid transmission (AAT)

An alternative potential mechanism for the lateral inhibition is that it could be driven by a poly-synaptic inhibitory pathway within the transverse pontine slice, for example recruiting recurrent connections with peri-LC GABAergic interneurons (as described by Breton-Provencher and Sur (2019)). To address this possibility, a series of LC recordings were made in the presence of a cocktail of pharmacological blockers of fast excitatory and inhibitory amino-acid transmission (AAT, comprising NMDA receptor antagonist D-APV (50μM), AMPA receptor antagonist CNQX (10μM), GABA_A_ receptor antagonist Gabazine (5μM), and Picrotoxin (50μM) to block both GABA_A_ and Glycine receptors). Following the previous chemogenetic LC activation protocol with PSEM308 (10μM) bolus at baseline followed by a wash-in of AAT antagonist cocktail with a second PSEM308 bolus. Following a 20 minute wash-out period, a final PSEM308 bolus was given (Figure S8A). AAT blockade resulted in a gradual increase in intracellular calcium across the recorded LC neurons (Figure S8A, 1.90±0.34 σ, n=36 cells). The overall increase in intracellular calcium signal following AAT blockade is potentially consistent with the blockade of ongoing tonic inhibition such as that from peri-LC GABAergic neurons, given that many of these neurons (located medial to the LC core) will be preserved in the transverse slices.

The magnitude of PSEM308 response after AAT blockade was compared for PSEM308 excited and inhibited cells. Although a significant difference was found between drug applications for inhibitory responses (n=10 cells, RM-ANOVA, p=0.0043), post-hoc testing revealing that this was not due to the effect of the antagonist cocktail on the inhibitory response but rather there was a decrease between the initial PSEM308 response and the final wash-out PSEM308 response (Figure S8B, Tukey HSD p=0.015). AAT blockade had no effect on the magnitude of the excitatory responses to PSEM308 (Figure S8C).

The lack of blockade of the lateral inhibition in the presence of a cocktail of AAT antagonists suggests that it is not dependent on recurrent inhibition via peri-LC GABAergic neurons, or indeed any other poly-synaptic pathway retained within the transverse pontine slice.

### Lateral inhibition occurs preferentially between spatially proximate neurons

To investigate the principle that the inhibitory effect required local release of noradrenaline from excited neurons the spatial organisation of the cells was explored within recording sessions. The hypothesis to be tested was that on average the inhibited neurons would be closer to an excited neuron than the population of unresponsive neurons (Figure 7). Within each slice, the distance between the centres of each inhibited LC neuron profiles and all PSEM308 excited neurons was measured, and similarly the nearest PSEM308 excited neighbour was found for all non-responsive LC neurons. Comparing the nearest neighbour distance for these pairs across the dataset showed that the distance from excited->inhibited cells (27.6±13.5μm, n=44) was significantly closer on average than excited->non-responder cells (37.6±19.5μm, n=47, two tailed t-test stat = −2.811, p value = 0.0061). Conversely, this was also not consistent with a picture of simple diffusion of NA-mediated “surround” inhibition as more than 30% of the non-responsive LC neurons were closer to an excited cell than the mean distance of the inhibited cells from the excited cells (within 27µm).

**Figure 7.**
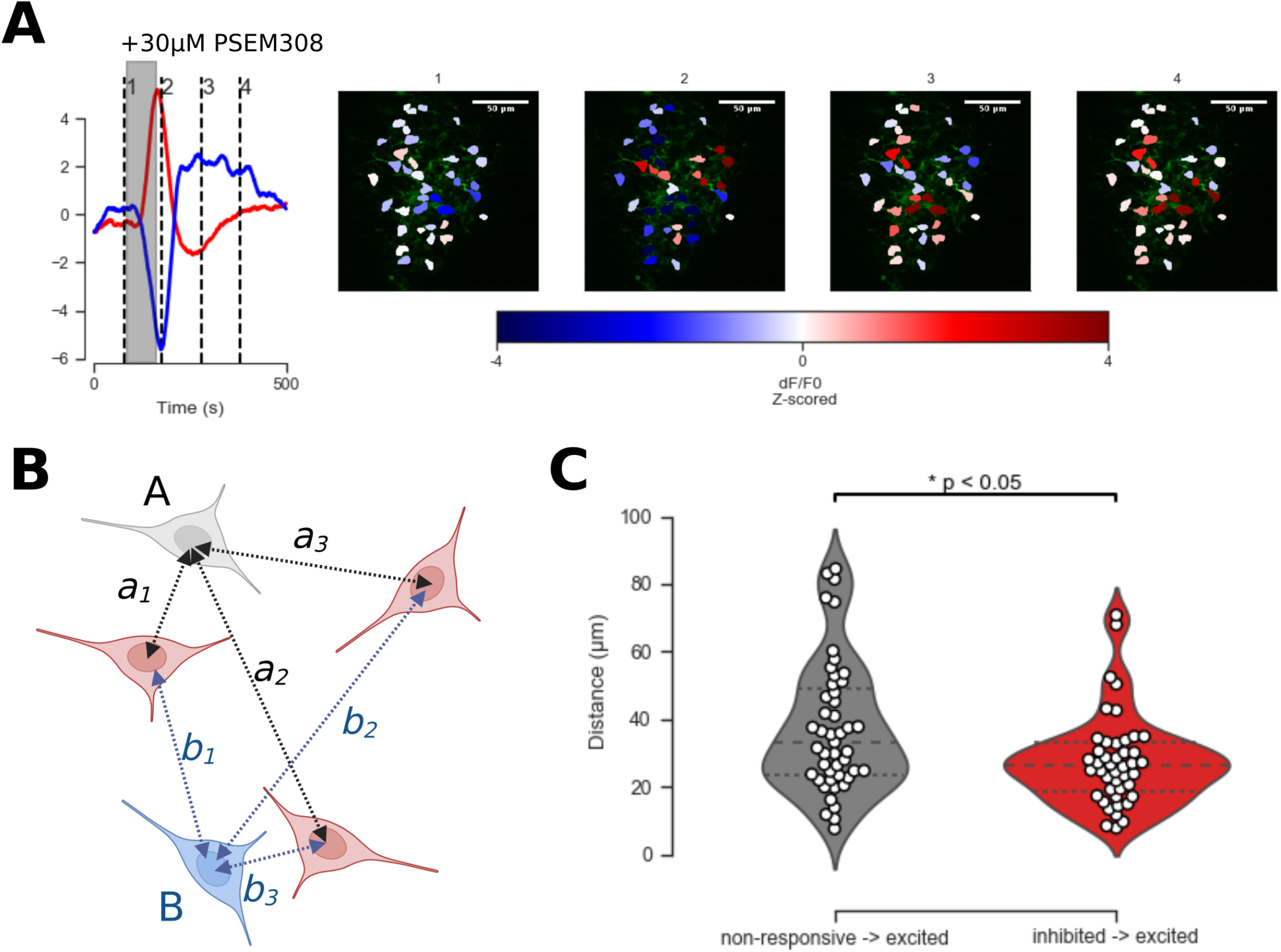
Spatial organisation of lateral inhibition. A – LC neurons that were either excited or inhibited following bolus PSEM308 perfusion (grey box, left panel) appeared to be clustered in space (subpanels 1-4 on the right show heatmaps of GCaMP6s signal in individual neurons at time points 1-4 (dashed vertical lines) as shown on the left panel) B – graphical representation of the distances between the centres of pairs of cells. For each PSEM308 non-responsive cell (A) and each PSEM308 inhibited cell (B), the distance to the nearest PSEM308 excited cell was found (in this case a1 & b3 respectively). C - Comparison of the distance between non-responsive→excited vs inhibited→excited nearest neighbour pairs. Inhibited→excited pairs were significantly closer together than non-responsive→excited pairs (n=47 non-responsive→excited pairs, n=44 inhibited→excited pairs, 9 slices from 5 animals, two tailed t-test stat = - 2.811, p value = 0.0061)

### Lateral inhibition is predominantly cross-modular

The CAV-PRS-GCaMP6s and CAV-PRS-PSAM vectors were retrogradely targeted to different LC modules to assess whether these inhibitory interactions were indeed a targeted, modular phenomenon, or rather reflected a generic property of LC neurons to provide an arbitrary inhibition to neighbouring cells. For these experiments CAV-PRS-GCaMP6s was injected into the olfactory bulb, chosen as it has a large, specific LC efferent projection (Schwarz *et al*., 2015; Kebschull *et al*., 2016). Conversely, CAV-PRS-PSAM was injected into the spino-medullary junction targeting the descending spinally projecting tracts of LC neurons which similarly have been reported to have both distinctive inputs and specific output projections (Schwarz *et al*., 2015). The objective was to generate slices in which PSAM activation would be restricted to the bulbospinally projecting LC neurons (LC_BS_) and evoked calcium responses recorded from only olfactory bulb projecting neurons (LC_OB_), referred to hereafter as ‘cross-modular’ interaction (Figure 8A). These modules were chosen due to the large anatomical separation of their efferent projections, meaning each vector injection would be far removed from the other, preventing ‘overlap’ of vector exposure at either target site.

**Figure 8.**
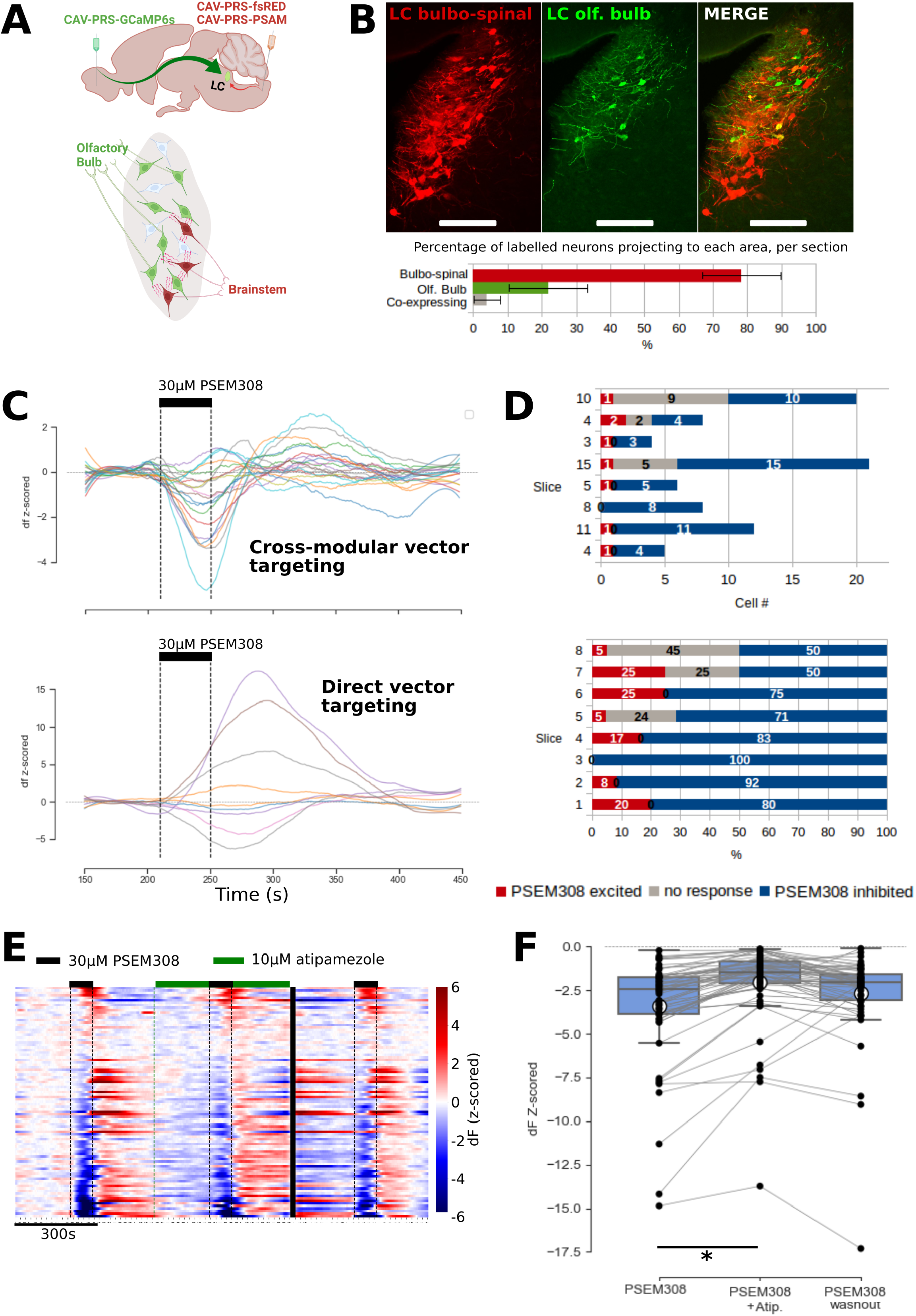
Cross-modular lateral inhibition of LC_OB_ by LC_BS_ neurons. A – Slices were prepared following retrograde targeting of two distinct LC modules with CAV-PRS-GCaMP6s injected into olfactory bulb and CAV-PRS-PSAM or CAV-PRS-fsRED into brainstem. B – This approach resulted in a higher number of transduced LC_BS_ neurons than LC_OB_ projecting neurons, and a subset of co-labelled cell bodies (n=3 sections, 1 rat, 60 cells). (For this assay CAV-PRS-fsRED was injected in the brainstem) C - Slices generated with cross-modular targeting showed predominantly inhibitory responses to PSEM308 in the LC_OB_ (top panel), compared to slices in which CAV-PRS-GCaMP6s and CAV-PRS-PSAM were co-injected (bottom panel). Plots show individual neuron level data (Savitsky-Golay filtered) from a single slice from each dataset. D - The predominance of inhibitory responses is highlighted further when examining the number (top panel) and proportion (bottom panel) of cells in each response category for each neuron per slice (n=8 slices, 4 rats) E – Raster heatmap plot of all neurons in this cross-modular dataset, sorted according to their response to PSEM308 (most excited at the top). The predominance of inhibition and the biphasic nature of responses are both evident. (n=93 neurons, 8 slices, 4 rats). The effect of atipamezole (10µM) attenuating the lateral inhibition is shown and which recovers after washing. F – Pooled data showing a significant block of the lateral inhibition by atipamezole (10µM). RM-ANOVA F stat =23.815, df=2, p-value < 0.00005, Tukey HSD * - P<0.05) (Boxes show median ± IQR, whiskers show 1.5x IQR, mean – white circles).

Immunohistological assay of sections taken from rats with CAV-PRS-GCaMP6s injected into the olfactory bulb and CAV-PRS-fsRED (Hayat *et al*., 2020) injected into the brainstem bulbospinal tracts, confirmed expression of both vectors in the LC (Figure 8B), in largely distinct cell groups with only a small degree of co-expression demonstrating some LC cells send divergent projections to both areas (4±3.8% co-labelling of total cell number n=3/60). In 2 photon recordings in slices with LC_OB-GCaMP6s_ neurons, and LC_BS-PSAM_ neurons, PSEM308 (30μM) mediated activation of the LC_BS_ module produced overwhelmingly inhibitory responses in the LC_OB_ module (60/93 neurons, 64.5%), with most of the remainder non-responsive (25/93 neurons, 26.9%) and a few excitatory responses (8/93 neurons, 8.6%). Representative PSEM308 responses for a slice generated with cross-modular vector targeting, compared to direct vector injection, illustrated the predominance of inhibitory responses following cross-modular activation (Figure 8C). This pattern is visible throughout the dataset of recordings (Figure 8D&E).

Atipamezole (10µM) was again used to test for the involvement of α2R signalling in this interaction (Figure 8E&F) and it had a significant effect on lateral inhibition of LC_OB_ neurons (RM-ANOVA, p<0.00005). The atipamezole blockade of α2Rs decreased the magnitude of lateral inhibition compared to the initial PSEM dose (from −3.41±2.98 σ to −2.06±2.31 σ, a decrease of 39.6%, Tukey HSD p=0.014). Atipamezole had no effect on the PSEM308 excitation in LC_BS_ /co-projecting neurons (RM-ANOVA F stat =0.9997, df=2, p=0.39). This suggests that the cross-modular lateral inhibition of LC_OB_ neurons, evoked by LC_BS_ activation, is mediated by α2R signalling.

To test whether this cross-modular inhibition was reciprocal, an additional set of recordings was made with the targets of each vector injection swapped - CAV-PRS- GCaMP6s targeting LC_BS_ neurons, and CAV-PRS-PSAM to target LC_OB_ neurons (Figure 9C and S9). This showed similar findings with a majority (76.5%) of the LC neurons exhibiting inhibitory responses to the 30μM PSEM308 bolus (13/17 cells) and a small number showing excitation (17%, 3/17) (Figure 9C). Lateral inhibition of LC_BS_ neurons was again significantly reduced by α2R blockade with atipamezole (RM-ANOVA, F stat =12.2667, df=2, p-value = 0.0002). Atipamezole decreased the magnitude of the inhibition (from −4.20±1.16 to −2.86±1.17 σ, by 31.9%, Tukey HSD adjusted P-value = 0.0175, Figure S9D). Atipamezole had no effect on the PSEM308 excitation (RM ANOVA, p=0.59).

**Figure 9.**
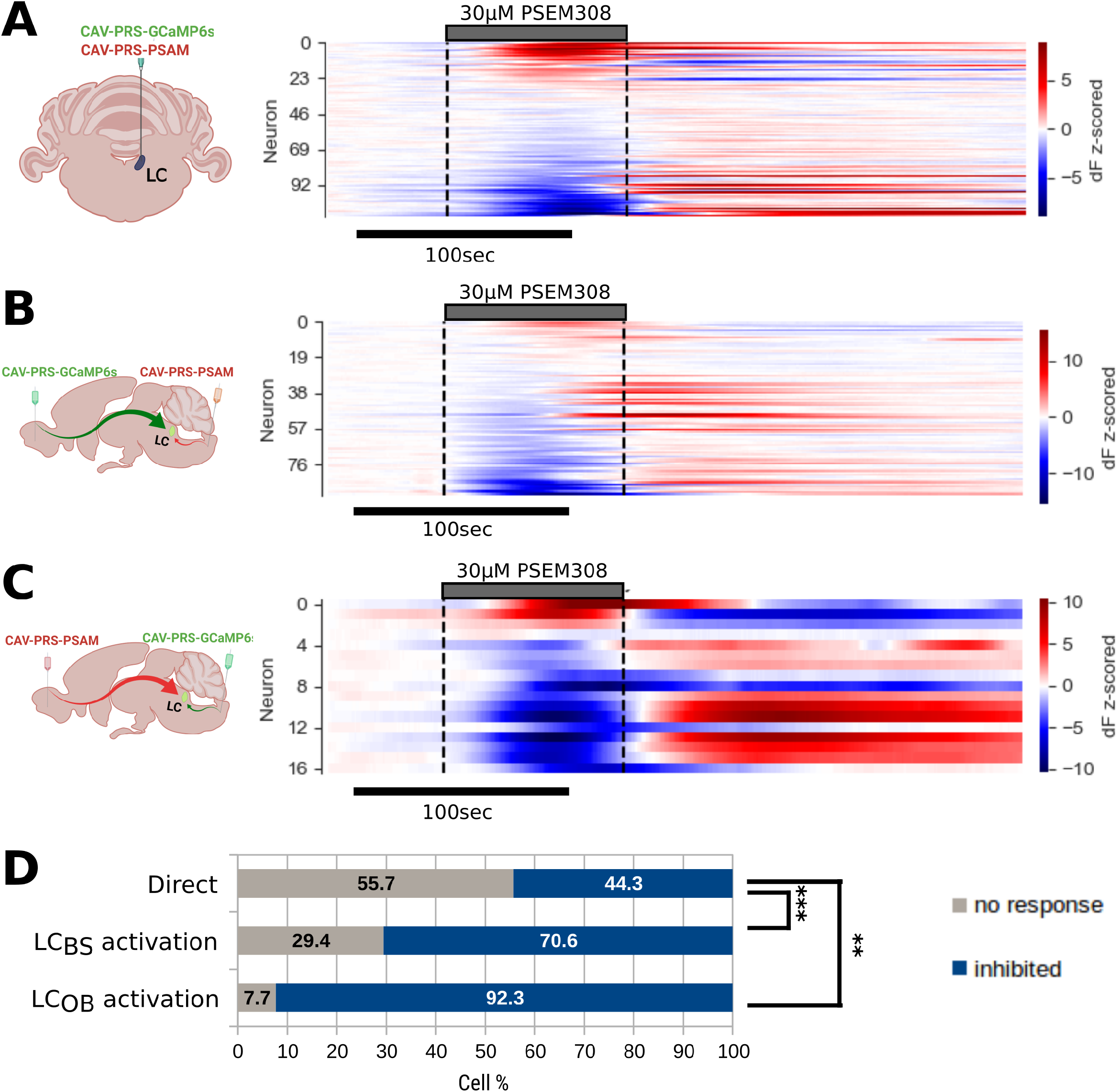
Comparing proportion of neurons in each response category for direct vs modular PSAM transduction. Raster plot of all neurons for each dataset sorted by response to 30μM PSEM308 A - direct LC transduction, B - LC_BS_ activation vs LC_OB_ imaged C -LC_OB_ activation, LC_BS_ imaged (2 slices from 1 rat) D – Differing transduction & activation strategies resulted in significant differences between proportions of inhibited vs non-responsive neurons following PSAM activation for both ‘cross-modular’ datasets compared to the direct injection dataset. Plot shows proportion of LC neurons in each category for ease of comparison, statistical testing was performed using Chi-squared test (excluding excited neurons) in figure 9 supplementary table 1 (Chi^2^ value = 18.6, p-value 9.2e-05) with FDR correction for pairwise comparisons, (** - p<0.01, *** - p < 0.001).

Lateral inhibition appeared to be more prevalent in slices with modular targeting (injections targeting LC_BS_ and LC_OB_ modules separately) compared to direct transduction of the LC with both CAV-PRS-GCaMP6 and CAV-PRS-PSAM (Figure 9). Significant differences were found between proportions of inhibited vs non-responsive neurons following PSAM activation for the different transduction strategies (Figure 9D*, Chi^2^ value = 18.6, p-value 9.2e^-05^*). (excitatory responses were excluded as they would be inherently different due to the different transduction strategies). Post-hoc testing revealed that compared to direct LC targeting of both vectors, cross-modular targeting resulted in a greater proportion of inhibitory responses, and fewer non-responsive neurons (FDR corrected for multiple comparisons: p=0.005 for direct vs LC_OB_ activation, corrected p=0.0025 for direct vs LC_BS_ activation). This demonstrates that specific modular activation preferentially evokes inhibitory responses in LC neurons belonging to other modules, rather than stochastic non-specific transduction after direct LC injection.

### Cross-modular inhibition generates contrast enhancement of synaptic drives

The effect of cross-modular inhibition on the activity responses to a synaptic drive was investigated by simulating pairs of modules (each with 3 neurons), with reciprocal α2R inhibition between modules (Figure S10A). Phasic activation of one module (M1) was modelled by delivering a 15Hz train of excitatory post-synaptic potentials, EPSPs lasting 3 seconds (Figure S10B). This drove the neurons in M1 to increase their rate of action potential discharge in a phasic-like burst (∼3Hz). As the strength of α2R inhibition increased, a cross-modular inhibition emerged affecting the spontaneous activity of the second module (M2). This manifested as first an increase in the interspike interval (ISI) followed by a complete silencing of the inhibited module (Figure S10C).

The relative change in activity across the modules produced by α2R inhibition was quantified by calculating a contrast enhancement index:

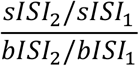

Where: *sISI*_1_ *sISI*_2_ = *ISI during the phasic stimulation period for module 1 & 2, respectively (for module 2, this was inclusive of the first spike immediately preceding, and following, the period of phasic activation)*. *bISI*_1_ *bISI*_2_= *ISI during the basal (pre-stimulus) period for module 1 & 2, respectively*

This contrast enhancement index increases dramatically as the strength of lateral inhibition is increased in the modelled populations, before plateauing as total cross-modular silencing occurs (Figure S10C). This mechanism therefore amplifies the differential in activity between the two modules.

### Lateral inhibition produces reciprocal rebound waves of activity in LC networks

Examination of the heat maps of LC activity following the chemogenetic modular activation and lateral inhibition (Figure 8E&F) also revealed that these were followed in many cells by a reciprocal change in the tonic activity that could last for 100-300s after the initial response. This was most clearly demonstrated in the neurons that showed initial lateral inhibition with subsequent rebound excitation (72.5% of cells, Figure 10A&B). This rebound excitatory drive could be powerful (4.9±5.6 σ, Figure 10C) and was also apparent in the directly transduced cells and in the single cell extracted dF/F_o_ time series (Fig 5A, Fig 8C). A rebound inhibition was also noted in some of the cells that were initially excited by PSEM308 but this was weaker and less common (17.4% of cells, Figure 8E & 9B, 10B&C). The periodicity of these biphasic responses was similar for the inhibition-excitation and the excitation-inhibition dyads with ∼70s between the peak of the initial response and the peak of the subsequent obverse effect giving a frequency of around 0.007Hz (Figure 10C).

**Figure 10.**
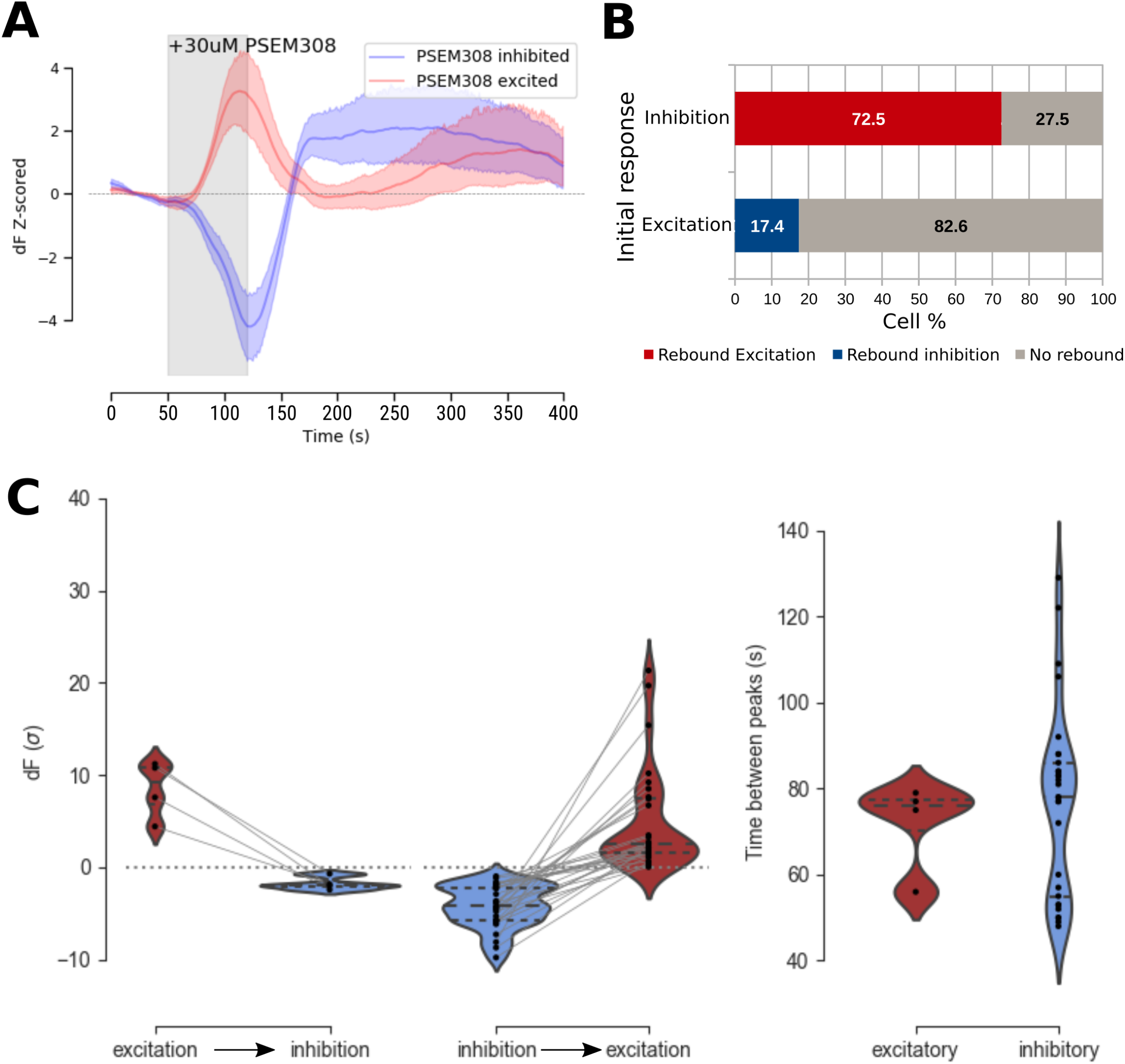
Biphasic responses following PSAM activation. A – Averaged time series data from a single slice showing PSEM308 excited and PSEM308 inhibitory responses, illustrates the biphasic nature of the inhibitory response, with the GCaMP6s signal diverging both during the initial PSEM308 response, and in the subsequent post-inhibition excitation. (mean +/- 95% C.I., n=11 excited neurons, n=16 inhibited neurons) B – Biphasic responses were more prevalent in neurons showing initial inhibition (29/40 neurons, 72.5%), than for neurons showing initial excitation (4/23 neurons, 17.4%) C – Left panels show the magnitude of initial and rebound responses for excitation- >inhibition and inhibition->excitation biphasic responses. Right panel shows the time between the peak of the initial response and rebound response for both excitation- >inhibition and inhibition->excitation biphasic responses. (Violin plot dashed lines show median and quartiles).

### Modelling biphasic responses with reciprocal α2R-inhibition and NA reuptake

The biphasic response properties of LC neurons with excitation followed by inhibition (for example by noxious or salient stimuli such as a foot pinch) are known to involve α2Rs, activated by the noradrenaline released following elevated firing and presumed to be a feedback autoinhibition. (Cedarbaum & Aghajanian, 1977; Aghajanian & VanderMaelen, 1982; Andrade & Aghajanian, 1984; Williams *et al*., 1985). In addition to biphasic responses following PSEM308 excitation, the evoked lateral inhibition observed in neighbouring cells also had a biphasic character (initial inhibitory response followed by rebound excitation, Figure 8E & 9). It was hypothesised that this rebound excitation in non-transduced neurons may be the result of suppressed firing of the PSAM-activated neurons, which normally provide ongoing lateral inhibitory tone. Simulations focused on replicating biphasic excitatory responses in ‘PSAM+’ neurons, and then on investigating whether concurrent biphasic responses (of initial inhibition followed by rebound excitation) could be observed in neighbouring neurons, which were not directly excited by PSAM currents.

With this model generating biphasic responses in terms of autoinhibition, the major aim of replicating biphasic responses in neighbouring neurons could be attempted. Paired simulations were implemented, with one ‘PSAM+’ neuron receiving the same transient increase in excitatory current as before, approximating chemogenetic activation. This PSAM+ neuron was connected via reciprocal α2R inhibition with a neighbouring neuron, which did not receive the excitatory ‘PSAM’ stimulus (Figure 11). α2R inhibition was implemented using the proposed method to model NET occupancy and its relationship to synaptic temporal dynamics (the sigmoid transfer function is shown in Figure 10B). In these simulations, both the direct PSAM excitation and the corresponding inhibition in the neighbouring neuron, is followed by a biphasic second component. In the PSAM excited neuron, there is a period of suppressed firing due to both autoinhibition, and lateral inhibition, augmented by an increase in firing rate of the neighbouring neuron as it is freed from ongoing inhibition (Figure 11, biphasic responses especially clear in the traces showing V m and those showing instantaneous spike rate for each neuron). As with simulations modelling autoinhibition, the prolonged duration of lateral inhibition was mediated by the prolonged decay kinetics of gα2 for spikes occurring during high frequency firing (Figure 11)

**Figure 11.**
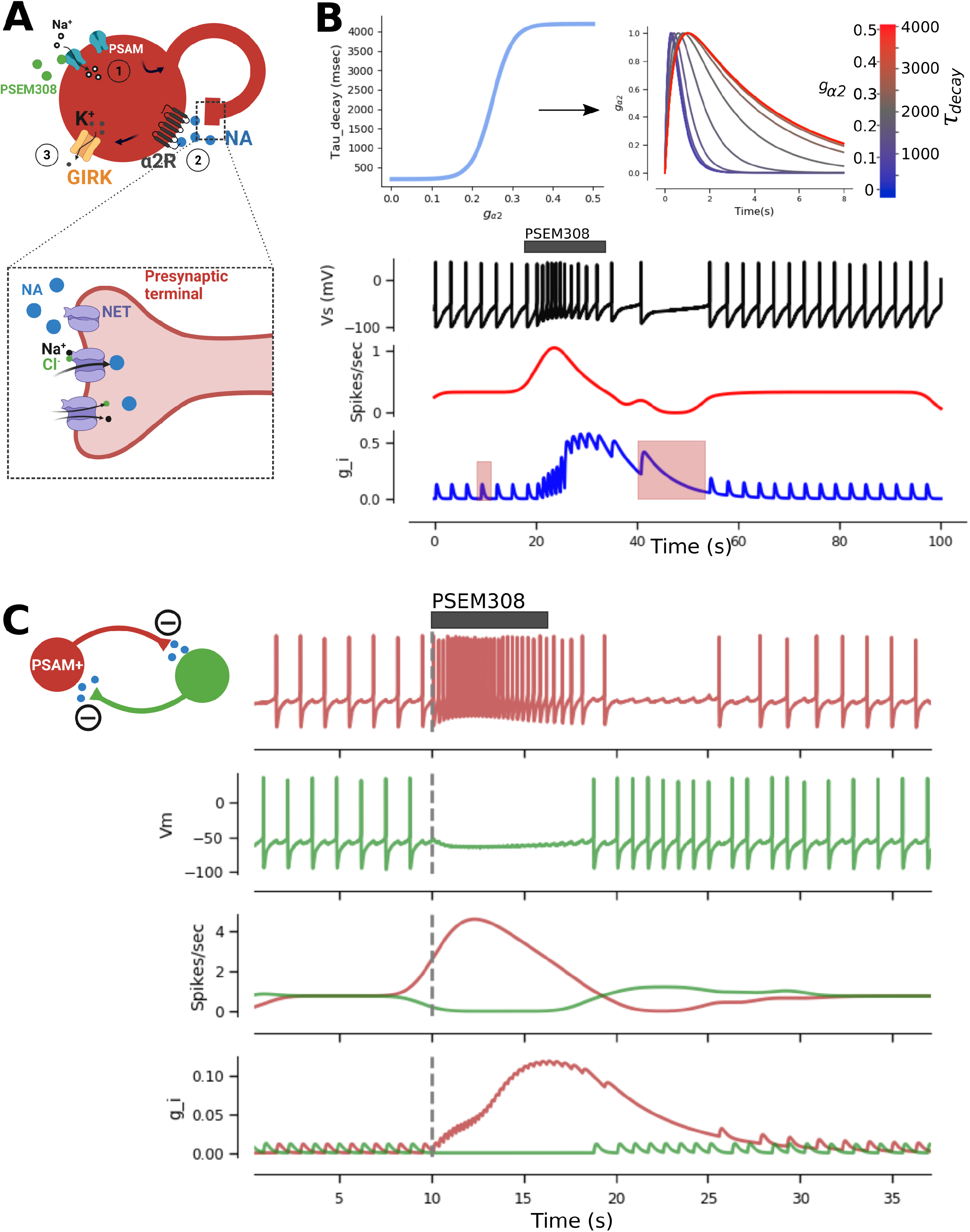
Modelling NA re-uptake saturation replicates biphasic auto-inhibition following PSAM activation in LC neurons. A – Schematic of modelling implementation. 1 - PSAM receptor activation is modeled as a transient increase in gPSAM, an excitatory sodium conductance, to drive depolarisation and increased firing rates. 2 – Noradrenergic release following action potentials is modeled as a transient increase in gα2, an inhibitory potassium conductance representing α2R activation and (3) subsequent GIRK activation. Inset panel shows NET reuptake of released NA, by cotransport with sodium and chloride ions. B – Saturation of NA re-uptake resulting from high NA release was modeled by use of a double exponential to describe spike evoked g α2 transients, with the decay constant (τdecay) of this exponential derived from the sum total of g α2 at the time of each spike, via a sigmoid transfer function representing ongoing NET occupancy.Top-left panel shows the sigmoid transfer function to derive the decay constant from g α2 (the sum of inhibitory α2 conductance in the spiking neuron, at the time of each spike). Top-right panel shows synaptic transients generated at different g α2. By modelling NA re-uptake saturation in this manner, the biphasic response of LC neurons to transient excitation can be replicated, as shown in the bottom panel traces. Following an increase in firing frequency due to the excitatory PSAM current, a post-excitation silencing can be observed as a drop in firing rate below basal frequencies. This post-activation inhibition is mediated by the increase in the decay time of spike-evoked g α2 transients during basal periods vs at times when NA release is already high (red boxes on g α2 plot, bottom panel, indicate the difference in exponential decay constant for release events occurring at basal firing vs following high-frequency activation). C – LC neuron models were set up in reciprocally connected (by α2R inhibition) pairs, in which one neuron would receive transient excitation replicating PSAM activation, and NET occupancy would be modelled as previously. Biphasic responses were observed for both directly PSAM-excited neurons, and paired neurons receiving lateral inhibition. The initial silencing of ongoing spiking in the neighbouring neuron is followed by a period of reversed responses, where the PSAM activated neuron is inhibited by auto-inhibition and the previously inhibited neuron is therefore freed from lateral inhibition. This facilitates an increase in firing frequency in the non-transduced neuron, further prolonging the inhibitory period in the PSAM activated neuron.

## Discussion

The locus coeruleus is organised into modules with groups of neurons innervating target fields with a degree of specificity (Foote *et al*., 1983; Loughlin *et al*., 1986; Howorth *et al*., 2009b; Kebschull *et al*., 2016; Hirschberg *et al*., 2017; Chandler *et al*., 2019; Poe *et al*., 2020). This functional projection architecture manifests in the LC but with a weak topographical organisation of somata with a degree of clustering but considerable intermingling. This anatomical organisation offers opportunities for co-ordinated interactions between groups of cells with related or contrasting functions. Imaging the network activity within the LC shows how the α2R-mediated inhibitory mechanism powerfully shapes the emergent patterns of activity. It is apparent that this α2R effect goes beyond feedback inhibition to be organised as a process of targeted lateral inhibition.

This lateral inhibition has many characteristics suggestive of vesicular synaptic release rather than the volume transmission model of LC neurotransmission (Zoli *et al*., 1998). It is relatively rapid for a G-protein coupled signalling event, exhibits frequency-dependent facilitation compatible with a calcium dependency and single action potentials can generate discrete postsynaptic events. The incomplete blockade by TTX suggests that the phenomenon is not only mediated by release from axon collaterals but is also mediated by somatodendritic release, as has been suggested by in vivo LC infusion of TTX with persisting release of noradrenaline as assayed by microdialysis (Singewald *et al*., 1994). Such a TTX-insensitive mechanism has also been supported by investigations in dissociated LC cells (Huang *et al*., 2007). The temporal kinetics of the noradrenergic signalling event are faster in the pontine slices (500-1500ms to the peak α2R-current) against a mean of 1820ms to onset of the noradrenaline release event in dissociated LC cells. The faster release kinetics agrees with the findings of Courtney and Ford (2014) and also the kinetics of noradrenaline release imaged using genetically encoded noradrenaline sensor GRAB_NE1m_ (Feng *et al*., 2019), perhaps reflecting the better-preserved anatomical arrangement with intact dendrites / spines and maintained co-localisation of calcium channels and vesicular release machinery.

The LC noradrenergic inhibition is distinct from the role played by the peri-LC GABAergic inhibition (Breton-Provencher & Sur, 2019) in terms of its slower onset and prolonged time-course. Both mechanisms are capable of regulating the tonic activity of LC neurons. This GABAergic mechanism has been reported to also be capable of providing feedforward inhibition of coherent “phasic” inputs to the LC (Breton-Provencher & Sur, 2019; Kuo *et al*., 2020). It is yet to be shown whether this GABAergic-shell inhibitory mechanism has any modular specificity or whether it can be recruited by activity within the LC itself (Breton-Provencher *et al*., 2021). Our GECI recordings confirm that the LC neurons *in vitro* are under a tonic GABA/Glycinergic inhibition as reported previously using single cell electrophysiological (Cherubini *et al*., 1988).

GECIs are beginning to be employed to study neuronal activity within the LC (Breton-Provencher *et al*., 2022; Dvorkin & Shea, 2022). This approach has not been used to study an inhibitory process such as that seen following activation of α2Rs although a decrease in fluorescence intensity after a phasic excitation associated with maternal pup retrieval has been noted (Dvorkin & Shea, 2022). The use of calcium imaging to study inhibition is becoming increasingly accepted in tonically active neuronal populations (Favre-Bulle *et al*., 2018; Gong *et al*., 2020; Vanwalleghem *et al*., 2020). The presence of spontaneous firing in LC neurons *in vitro* slices means that there is an elevated level of GCaMP6s fluorescence that can be decreased by block of action potentials (TTX) or by application of an α2R-agonist (clonidine). This has enabled the temporospatial properties and network distribution of these α2R inhibitory actions to be mapped onto specific groups of LC neurons.

The spatial organisation of the inhibitory effects is noteworthy as it indicates that there is both a proximity element and a specific targeting of the interaction. Any LC cell has an increased probability of being inhibited if it is within a ∼30µm radius from the centre of an excited LC neuron. This spatial distribution is, however, also overlapping with the non-responsive cells and so the picture is not simply one of surround / bystander inhibition with a shell of inhibition around each directly chemogenetically-excited neuron (Zoli *et al*., 1998; Baral *et al*., 2022). The distance between some of the inhibited cells and their nearest excited neighbour (between 40 and 75um) also means that diffusion distances are too long for any bulk release of noradrenaline to be plausible (the diffusion distance of Noradrenaline was previously estimated to be ∼ 3µm from the release site (Courtney & Ford, 2014)) given its known short half-life and the active reuptake mechanisms. Also, the relative strength of the inhibition and the speed of onset also argue against a long-range diffusion. There is anatomical evidence for specific synaptic contacts between dendrites of noradrenergic neurons within the LC (Shimizu & Imamoto, 1970; Groves & Wilson, 1980) that could mediate these targeted local interactions. However, it is also conceivable that local diffusion from dendritic vesicular release sites close to, but not synaptically connected to, the target neurons could equally explain these results which have striking similarities to the kinetic reported by Courtney and Ford (2014). The use of genetically encoded noradrenaline sensors such as GRAB_NE1m_ may enable resolution of this longstanding issue if they are selectively expressed in LC modules (Feng *et al*., 2019).

The cross-modular α2R-mediated inhibition was strikingly uniform and more prevalent (70-90% of cells inhibited in opponent modules) than the lateral inhibition seen with stochastic transduction of LC neurons (44% inhibited) after direct injection of CAV. The push-pull cross-modular interaction, shown here between the rostrally projecting LC_OB_ neurons and the caudally projecting LC_BS_ neurons, means that an active module can suppress other modules and this in turn would tend to release the active module from incoming lateral inhibition (which exerts a tonic influence on LC activity as a whole). This has the immediate temporal effect of increasing the contrast in noradrenergic outflow to different CNS regions, potentially to suit the behavioural context, and may help to overcome the challenge of discretely altering the output of this tonically active nucleus with relatively diffuse synaptic inputs (Schwarz *et al*., 2015; Breton-Provencher *et al*., 2022). This push-pull may be engaged in the balancing up of whether to inhibit and thus deprioritise a noxious stimulus when circumstances demand (strong activity in the LC_BS_ module suppressing ascending LC modules that promote attention) or the converse where the sensory input is not gated by descending control from the LC and the cortical networks are engaged by the dominant ascending LC modules to assess its salience and context to motivate behaviour change (pain). Whether this reciprocal inhibition is generalisable to all the LC cross-modular interactions or whether it exhibits a behaviourally consonant specificity in its organisation is an important topic (as has been suggested by module specific differences in the functional responses to α2R -agonists (Wagner-Altendorf *et al*., 2019)). This will require further investigation with *in vivo* paradigms employing modality-specific stimuli to test for these hypothesised interactions across modules within the LC.

An additional network property that was observed is a rebound ‘flip-flop’ in activity that saw groups of neurons that were initially inhibited flip to an excited state. This is an emergent property of the LC network models with a likely dependency upon NET function and its prevalence across experimental settings suggests that it may be a fundamental mechanism within the nucleus. This rebound excitation produced a corresponding inhibition in the opponent module. This activity oscillation had a relatively slow time-course (frequency ∼0.007 Hz) and this may be related to the infra-slow oscillation that has previously been observed in the LC *in vivo* (Totah *et al*., 2018). Such infra-slow LC oscillations have recently been linked to the microarchitecture of non-REM sleep with a periodicity of around 50-70s (Matosevich & Nir, 2021; Osorio-Forero *et al*., 2021; Kjaerby *et al*., 2022; Morici & Girardeau, 2022) and the causal involvement of LC-prefrontal and LC-thalamic projections which operate in synchrony. It may be that the opponency between these two and the remaining LC modules may account for these oscillations. Beyond this specific sleep context this may represent a means to restore behavioural “vigilance” across modalities that were previously suppressed in a phase of attentional focus.

The presence of lateral and cross-modular inhibition has implications for the study of LC activity using opto- and chemo-genetic methods. While these allow an unprecedented level of control over the LC neuronal activity during behaviour it has not been widely appreciated that the activation of a subset of neurons may produce inhibition of another locally un-transduced population (see Hirschberg et al (2017)). This inhibition is powerful and can also produce a temporal tail of altered activity in the obverse direction. This may help to account for some of the unexpected observations in the field such as in the seminal early observations of Carter et al (2010) where optoactivation of the LC led to an apparent decrease in cortical noradrenaline release (that they linked to depletion of noradrenaline in terminals). Similar considerations are likely to apply to inhibitory opto-/chemo-genetic interventions and although this is better appreciated in terms of rebound firing in the transduced neurons (Hayat *et al*., 2020; Kjaerby *et al*., 2022) the consequences of release from tonic inhibition of other LC neurons should be factored into the analysis models. This confound to interpretation may be minimised by retrograde targeting of genetically encoded actuators to specific modules whereby any cross-modular inhibition is likely to resemble the underlying physiology.

There is a striking non-linear facilitation of the α2R-mediated inhibition with increasing presynaptic firing frequency. This likely reflects the known presynaptic calcium dynamics and was well recapitulated by including a standard model for calcium-dependent vesicle release (Nadkarni *et al*., 2010). This frequency-dependent augmentation indicates that shifts in tonic firing rates in the LC that are typically seen in vivo (say from 2-4Hz) will be associated with relatively small changes in the level of noradrenaline and local inhibition within the LC. However, phasic bursts where the LC neurons can reach instantaneous firing frequencies above 20Hz is likely to optimally engage the α2R-inhibitory mechanism. This will mean that afferent synaptic barrages that signal salient information in a particular sensory domain (i.e. olfactory) arriving at one LC module (LC_OB_) are likely to be effectively enhanced within the associated cortical territories and prioritised in contrast to the innervation targets of other LC modules. This likely constitutes a fundamental mechanism for differential actions of this ancient but polychromatic neuromodulatory system.

## Supplementary Material

**Figure S1.**
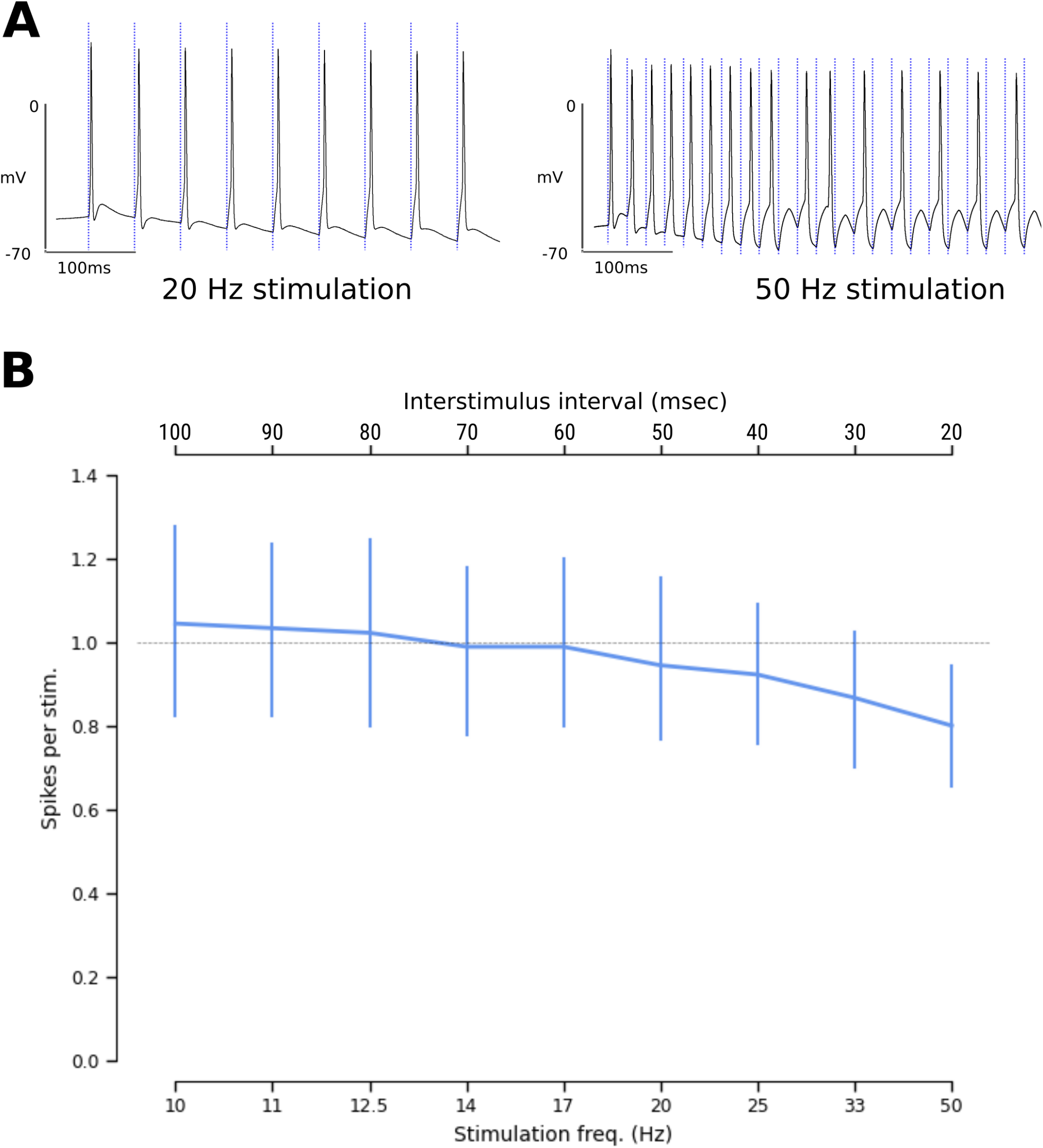
Optogenetic entrainment of firing in ChR2-transduced LC neurons. A – Light pulses were capable of entraining 1:1 firing at frequencies up to 20Hz (left panel) with accommodation occurring in some neurons at higher frequencies (50Hz). Blue dashed vertical lines indicate 10ms light pulses. B – Pooled data showing that optoactivation of ChR2-transduced LC neurons was reliable up to 20Hz and then showed a degree of accommodation when driven above this frequency (stimulus protocol was a 10 pulse light train, blue line indicates mean ± SD, n=9)

**Figure S2.**
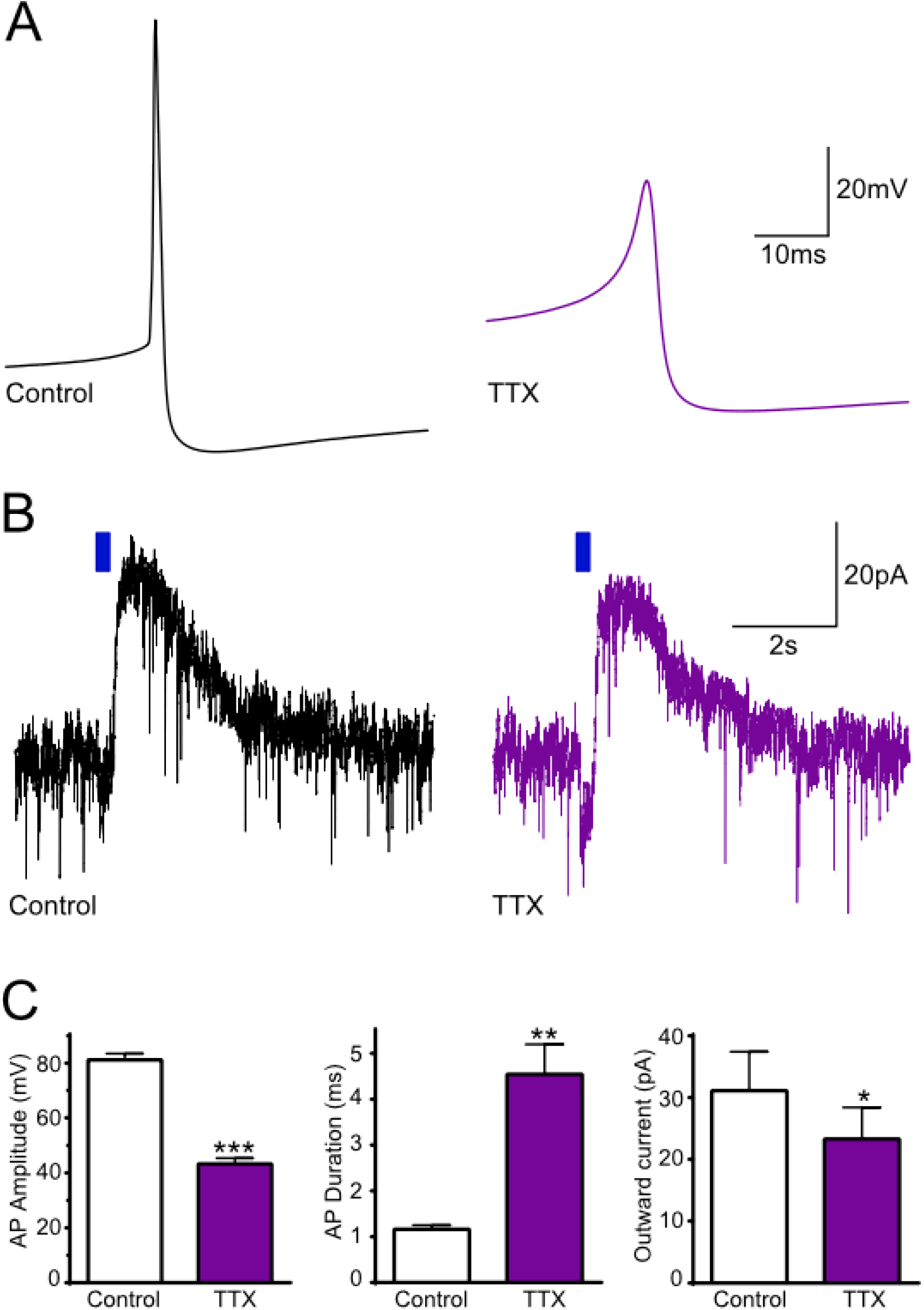
Evoked slow inhibition is mediated by a potassium current. A – Optogenetically-evoked outwards currents in recordings made using K-Gluconate based internal solutions remained stable for periods >15 minutes (20Hz x 1s 30ms light pulses). In contrast whole recordings made with a CsSO_4_-TEA internal solution showed a progressive decrease in outward current amplitude. Top panel shows representative outward currents at different time points from a single Cs-TEA recording, with blue dashed lines indicating opto-stimulation. (plots show mean ± standard deviation, n=5, 2 way RM ANOVA, F(1,8)=42.7, Sidak’s post hoc K-Glu vs Cs-TEA ** P<0.01, **** P<0.0001). B - The IV relationship of the inhibition was investigated by holding LC neurons at - 50mV with voltage steps (200ms) ranging from −40mV to −150mV either in control condition or at the peak of the light evoked outward current (200ms after the final LED pulse in a 20Hz x 1s 30ms light pulse train). The inhibitory current at each holding potential was obtained by subtraction. The current reversed at −110mV and showed evidence of inward rectification, in line with a potassium-mediated current

**Figure S3.**
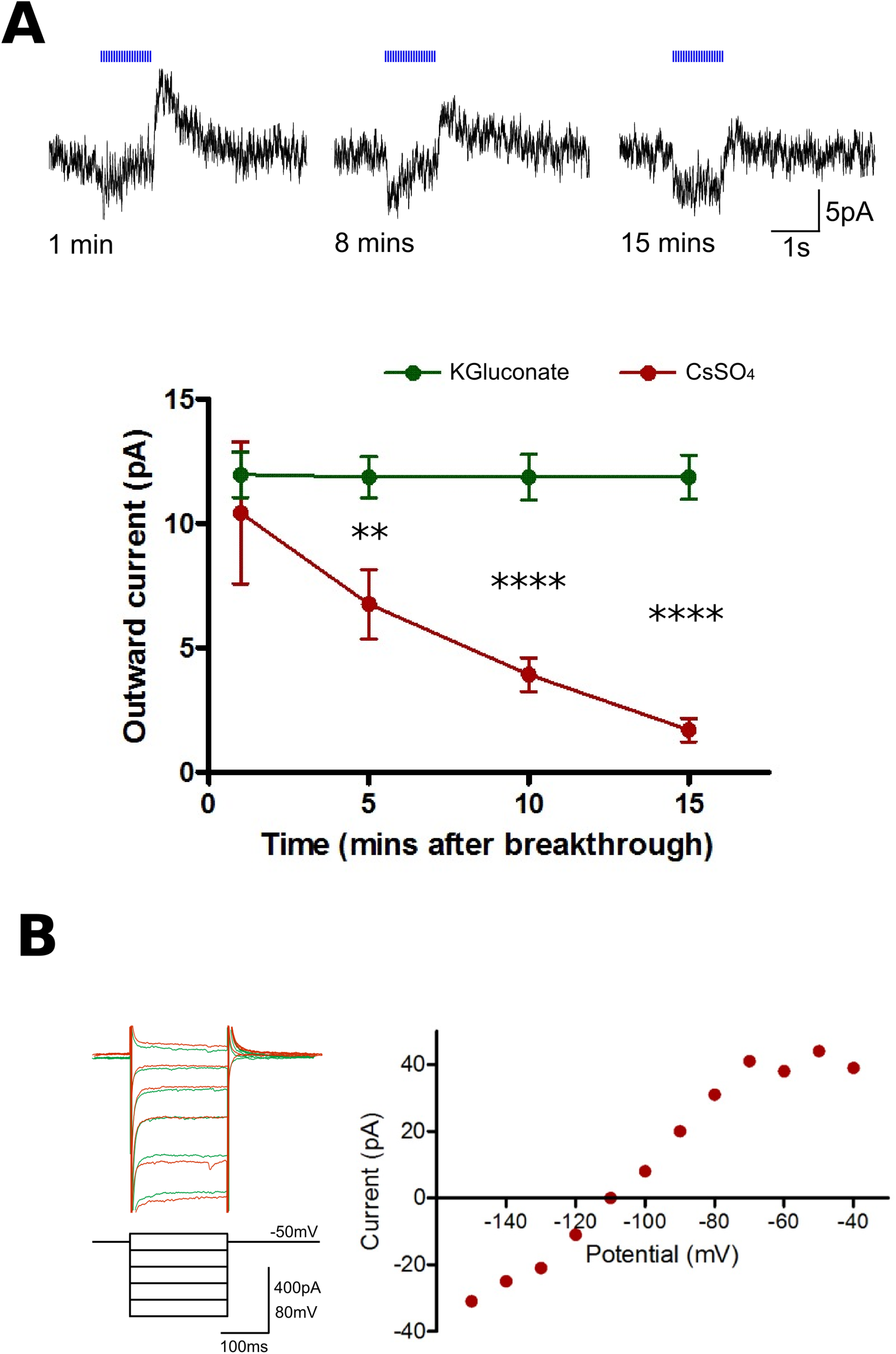
α2R-IPSCs are not dependent on fast sodium action potentials. A – TTX (500nM) abolished fast sodium action potentials in LC neurons, which subsequently fired spikes with a distinctively slower waveform with reduced amplitude. B – Optostimulation evoked α2R-IPSCs were attenuated but not abolished in the presence of TTX. C – TTX-resistant action potentials had significantly reduced amplitude (left panel) and significantly increased duration (middle panel) compared to normal sodium action potentials. α2R-IPSCs were significantly reduced in amplitude, but not abolished, in the presence of TTX. (Paired t-tests, n=5, * - p<0.05, ** - p<0.01, *** - p<0.001)

**Figure S4.**
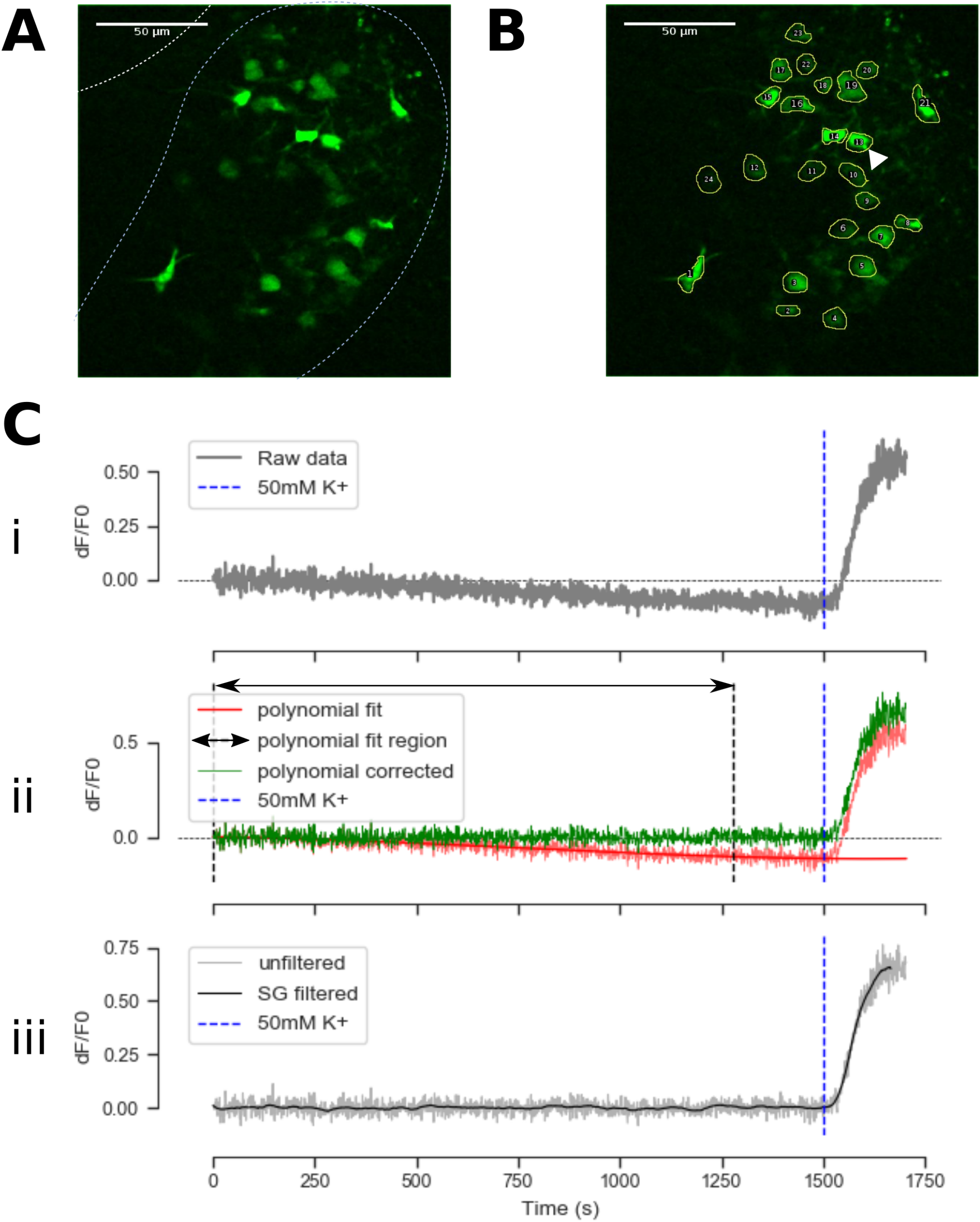
GCaMP6s time series pre-processing. A - GCaMP6s imaging was performed in ex-vivo transverse pontine slices containing the LC, with recordings of basal activity followed by a high potassium aCSF (50mM). GCaMP6s fluorescence was visible using two-photon excitation under basal conditions for the majority of cells, indicating that LC neuron ‘pacemaker’ firing may result in sufficient intracellular calcium levels at rest to produce visible GCaMP6s signal. Image shows a time-lapse Z-projection of basal fluorescence in a slice (over initial 200 seconds of recording). White dashed lines indicate the approximate boundary of the anatomical LC, and the boundary of the 4^th^ ventricle. B – Segmentation of individual GCaMP6s expressing LC neurons allowing extraction of time-series data. Segmented neuronal profiles are shown by yellow lines, white arrow indicates the neuron from which the traces in panel C were derived. Ci – Application of high potassium aCSF (50mM, onset at blue dashed vertical line) produced a striking increase in GCaMP6s fluorescence. Slow changes in GCaMP6s signal were also observed, which could represent either z-plane drift of the slice relative to the focal plane, or bleaching of GCaMP6s fluorescence over time in the case of gradual decay in signal. ii – Such slow drifts were corrected by fitting and subtracting a 3^rd^ order polynomial to the basal portion of the recording. iii – For measurement of transient responses, GCaMP6s timeseries data was processed using a Savitsky-Golay filter.

**Figure S5.**
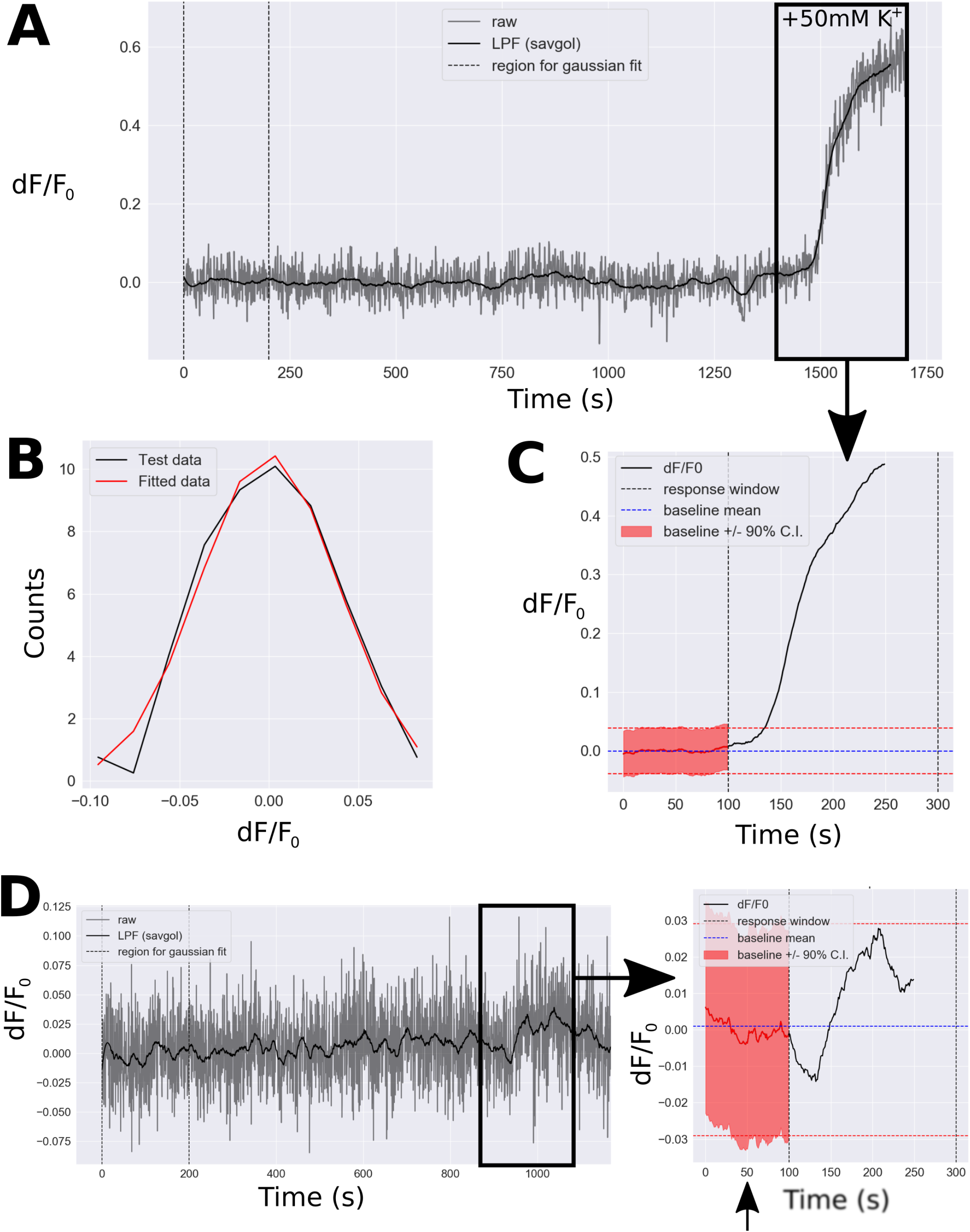
Detection of significant GECI responses. A – 2-photon recordings finished with a 50mM K^+^ aCSF wash (period indicated by black box), to identify healthy GCaMP6s expressing cells (as compared to dead cells or autofluorescent objects). Raw (unfiltered) data is shown, overlaid by the low pass filtered (Savitsky-Golay method) trace used for transient detection. B – A Gaussian model was fitted to the first 200 seconds of baseline imaging data for each cell, to estimate confidence intervals for detection of significant responses. C – Algorithmic detection of an excitatory response to high potassium aCSF, defined as the dF/F_0_ trace exceeding the 90% C.I. (centred around the baseline fluorescence value for the preceding 50 seconds) for at least 30 seconds, within a time frame of 50-250 seconds following the entry of high potassium aCSF into the perfusion system (shown by black arrow). D – In another cell from the same recording, this same method showed no significant response to high potassium aCSF (period indicated by black box). In the left panel, raw (unfiltered) data is shown, overlaid by the low pass filtered (Savitsky-Golay method) trace used for transient detection in the right panel. The black arrow indicates the time of high potassium aCSF entry into the perfusion system.

**Figure S6.**
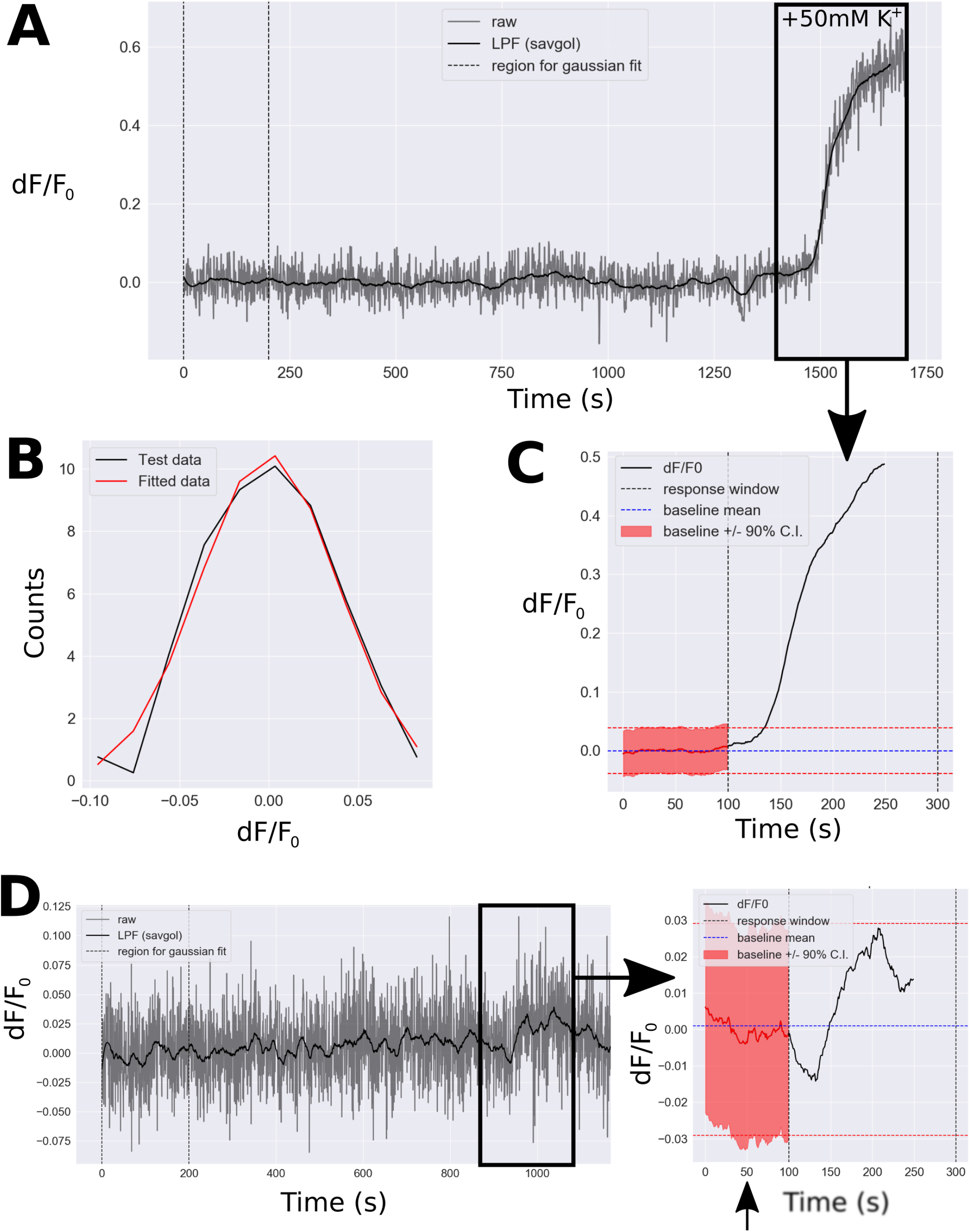
Detection of fast transient GECI events. A – Polynomial drift correction of imaging data, and calculation of the rolling average fSmooth baseline. Annotated orange regions show time and concentration of PSEM308 bolus. B – Gaussian model fit to first 200 seconds of imaging session, to allow quantification of confidence intervals for detection of significant transients. C – Gaussian noise model detection of an excitatory response, in this case to 30μM PSEM308. The fluorescence signal for this neuron exceeded the upper 90% confidence interval around the fSmooth baseline (derived from the Gaussian noise model fitted in S5B) for at least 30 seconds, within a time frame of 40-160 seconds following the entry of PSEM308 into the perfusion system, therefore an excitatory response was recorded. D – In another neuron, no significant response to 30µM PSEM308 was identified using this method.

**Figure S7.**
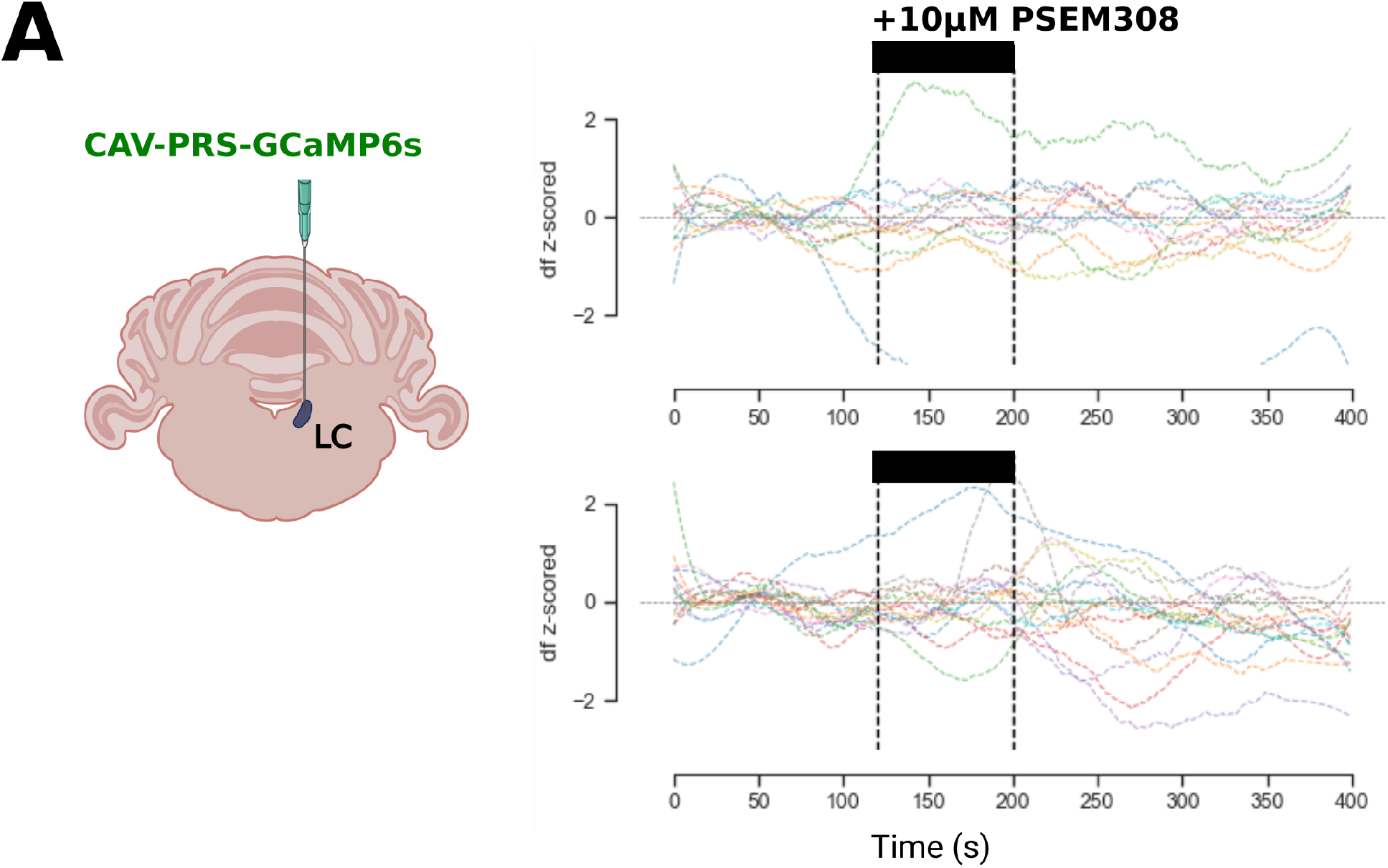
GCaMP6s expressing LC neurons do not respond to PSEM308. A – Plots of PSEM308 perfusion in 2 slices from a rat injected with CAV-PRS-GCaMP6s targeting the LC. In both plots, the large majority of GCaMP6s signals remain stable, with larger dF changes occurring in only a small number of cells, without temporal alignment to the PSEM308 bolus or other fluctuating cells (n=13 top panel, n=16 bottom panel)

**Figure S8.**
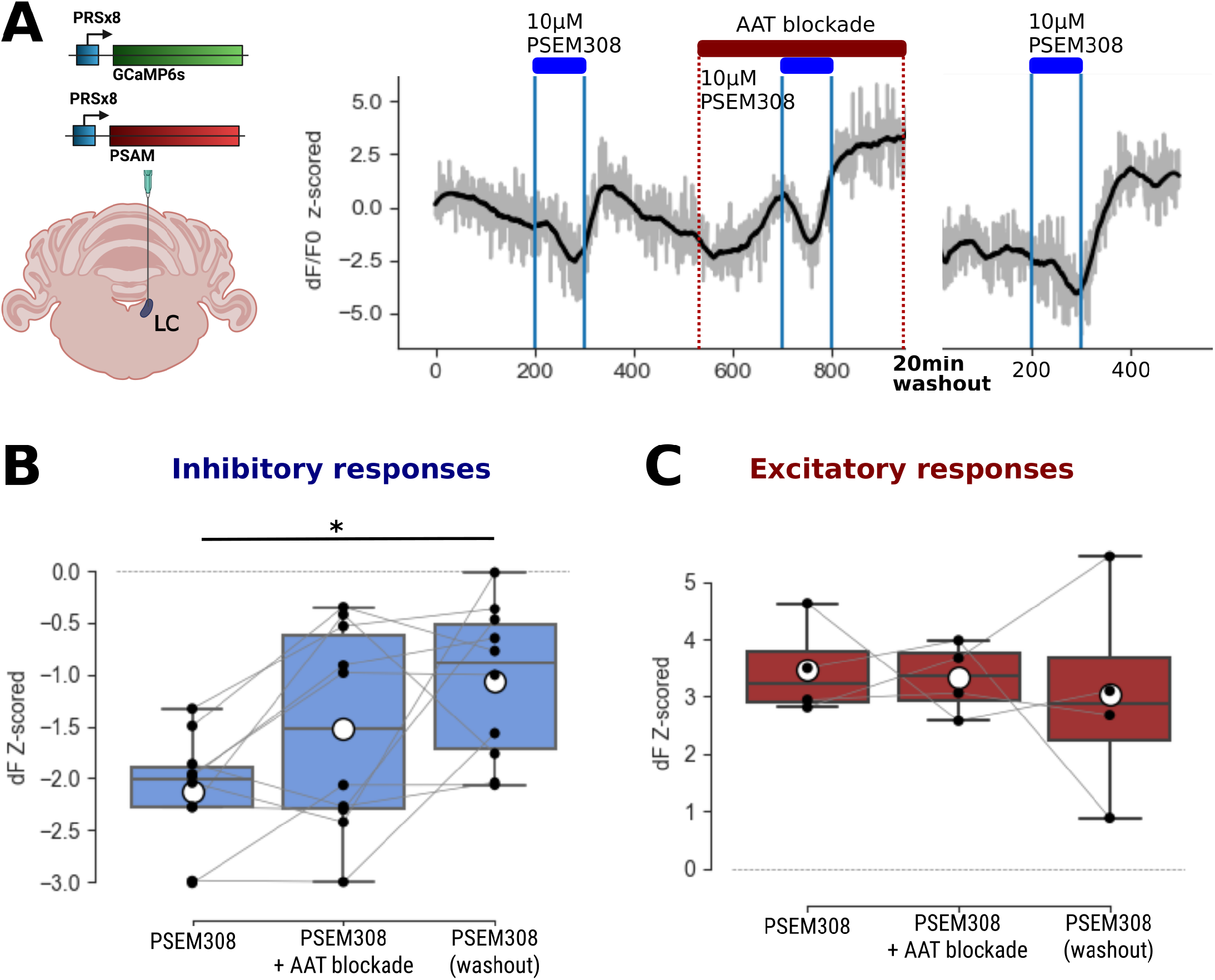
Lateral inhibition is insensitive AAT blockade with D-APV, CNQX, Picrotoxin and Gabazine. A – Slice recordings were made following direct transduction of LC with CAV-PRS-GCaMP6s and CAV-PRS-PSAM. Slices were perfused with a bolus of 10μM PSEM308 with/without AAT block, then allowed to recover for a 20 minute washout period. Right panel shows time series data for a single neuron whose lateral inhibition was not sensitive to AAT blockade. B & C – AAT blockade suggested a significant change in inhibitory (B, n=10 neurons, RM-ANOVA F stat = 7.474, df=2, p-value =0.0043, 3 slices, 1 rat) but not excitatory (C, n=4 neurons, RM ANOVA F stat = 0.111, df=2, p-value =0.9, 3 slices, 1 rat) responses following 10μM PSEM308. The only significant difference found was a decrease in inhibition between the initial inhibitory response to PSEM308 and the final inhibitory response to PSEM308 following AAT block washout (Tukey HSD adjusted P-value = 0.015). (Boxes show median + IQR, whiskers show 1.5x IQR).

**Figure S9.**
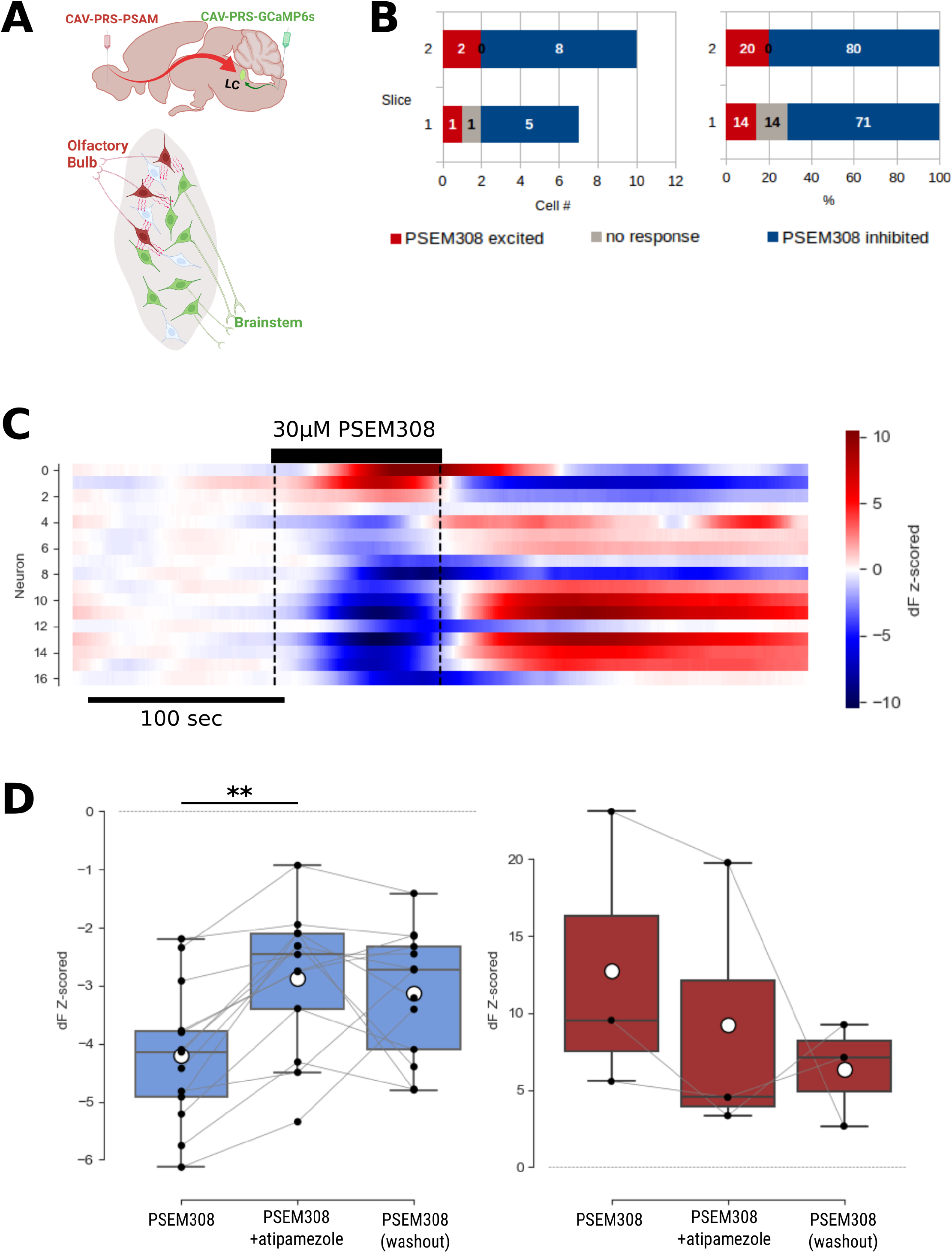
Cross-modular lateral inhibition of LC_BS_ by. A – Slices were prepared following retrograde targeting of two distinct LC modules – CAV-PRS-GCaMP6s targeting LC_BS_ and CAV-PRS-PSAM targeting LC_OB_. B – GCaMP6s responses of LC_BS_ neurons to PSEM308 activation of LC_OB_ neurons were largely inhibitory - left panel shows number of cells and right panel shows percentage of cells per slice in each response category. C – Raster heathmap plot showing PSEM308 responses for all neurons in this dataset, sorted according to their response from most excited top. (n=17 neurons, 2 slices, 1 rat). D – Pooled data showing the significant blockade by atipamezole (10µM) of the lateral inhibition (RM-ANOVA F stat =12.2667, df=2, p-value = 0.0002, Tukey HSD ** - p<0.01) but having no effect on the excitation (RM ANOVA F stat=0.5947, df=2, p=0.59). *(Boxes show median ± IQR, whiskers show 1.5x IQR, mean – white circles).*

**Figure S10.**
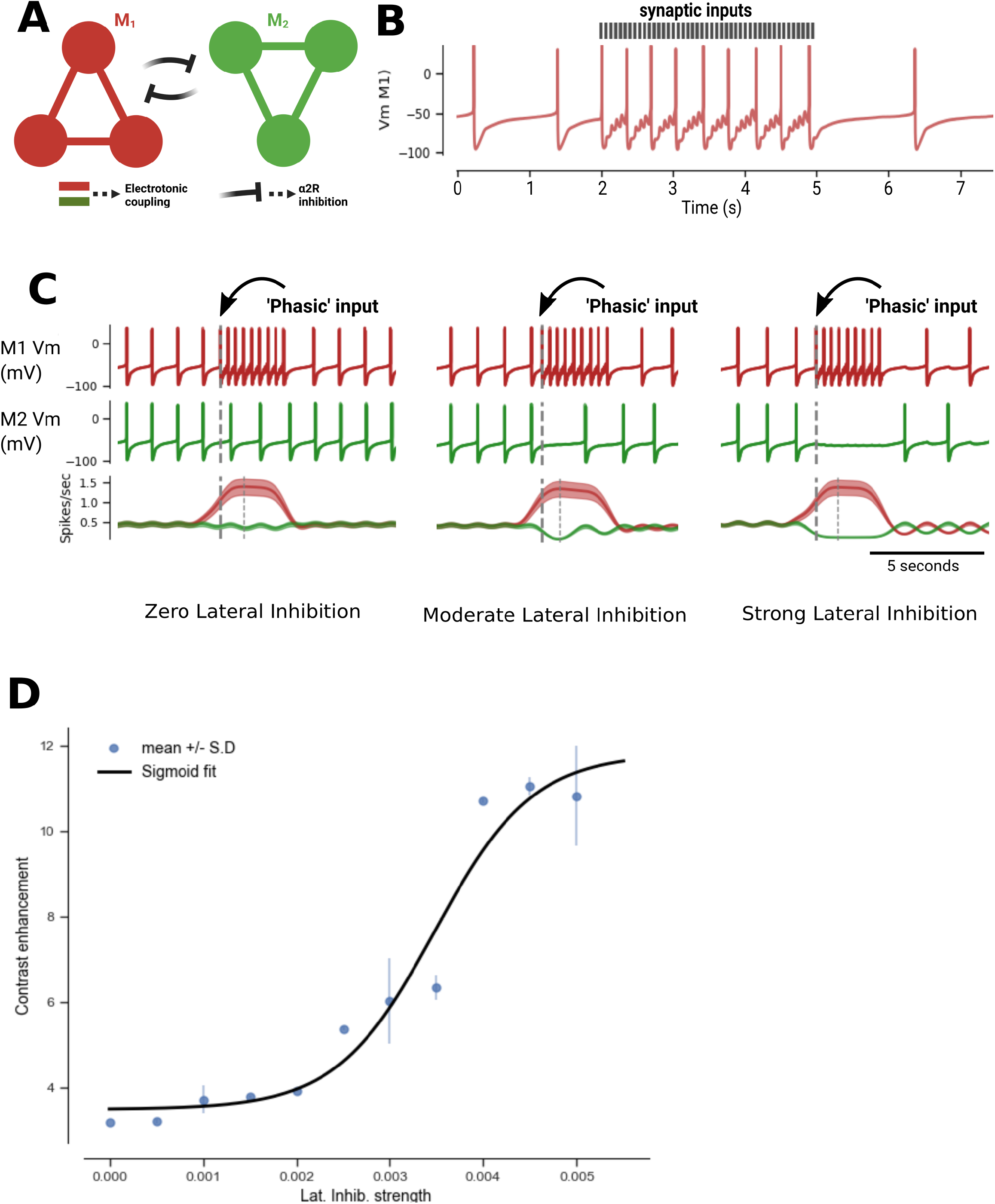
Lateral inhibition enhances contrast between LC modules during specific modular recruitment. A – LC neurons were simulated in a modular architecture, as populations of 6 neurons arranged into two reciprocally inhibitory modules, with intra-modular coupling via gap-junctions B - ‘Phasic’ activation of one module was simulated by delivery of a 15Hz, 3 second train of synaptic inputs, calibrated to drive action potential firing when the recipient neuron was close to threshold C – Example simulations with varying strength of lateral inhibition (none, moderate, and strong) show that the potency of cross modular silencing increases with stronger lateral inhibitory connections. (Vertical dashed grey lines show phasic stimulation onset (thick line). D – Contrast enhancement increases sharply alongside the strength of lateral inhibition, before plateauing as total modular silencing occurs. Blue circles and error bars show contrast enhancement index calculated from simulated data (mean +/- S.D.), and black line shows fitted sigmoid curve

## Appendix 1

The model consists of two compartments (soma and dendrite), in which the rate of membrane voltage is described by differential equations which depend on the sum of ionic conductances (all currents are given in µA/cm^2^ of membrane). For a given neuron *i* :

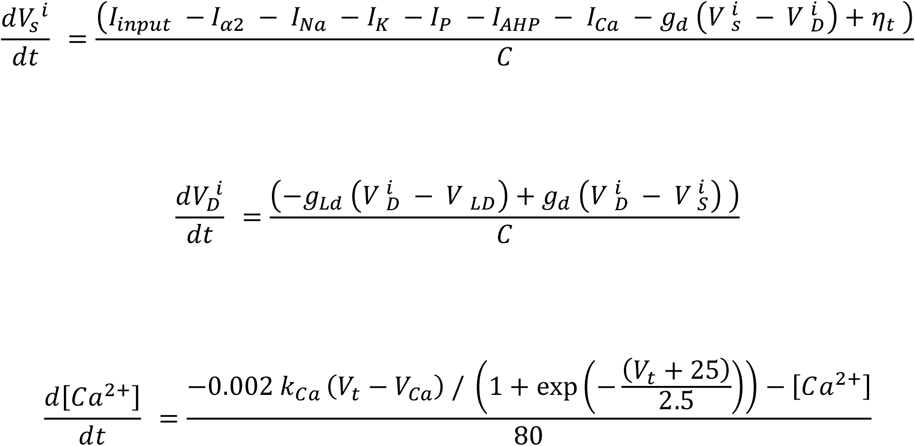

Where:

*V_s_* = somatic membrane voltage *V_D_ =* dendritic membrane voltage

*g_d_* = somato-dendritic conductance *g_Ld_*= dendritic leak conductance *[Ca^2+^] =* internal *Ca^2+^* concentration

*V_Ca_* = reversal potential for Ca current *I_input_*= experimentally applied intracellular current

*η_t_* = white noise function (mean=zero, scale=0.1) approximating stochastic influences

***n.b.*** Intracellular calcium concentration was modeled in a voltage dependent manner, approximating voltage gated calcium influx into the neuron without explicitly modelling these conductances in the Hodgkin-Huxley formalism.

The equations governing individual ionic currents were:

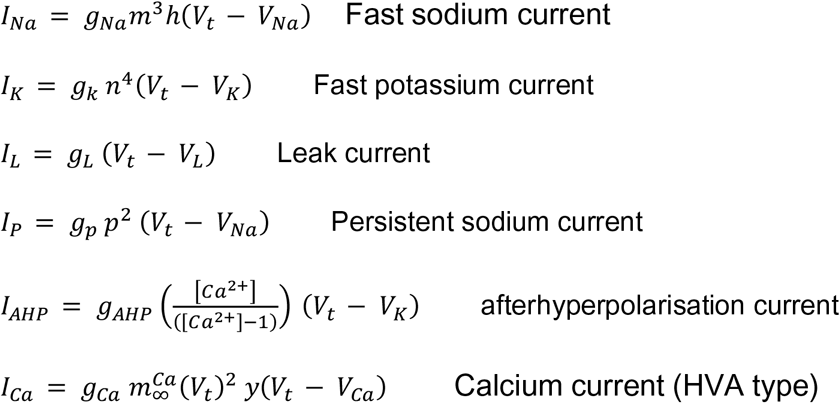

The fast sodium current activation variable *m* was assumed to be fast, and replaced by the asymptotic value:

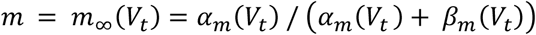

where:

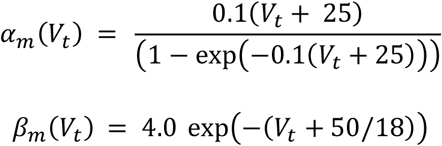

The gating variables *h, n,* and *p* obeyed the following differential equations:

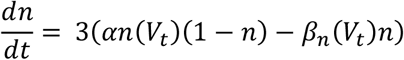

Where:

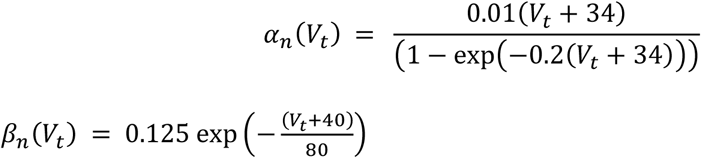

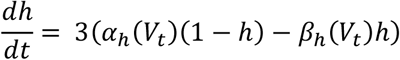

Where:

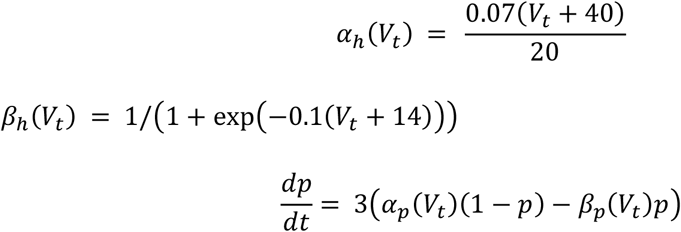

Where:

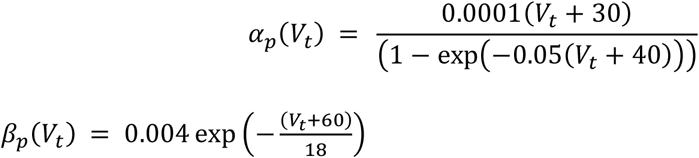

The calcium current activation variables were governed as follows:

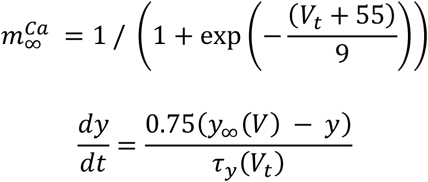

Where:

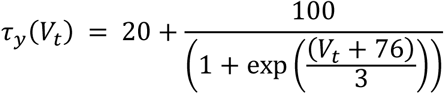

### Model parameter values

| Parameter | Value | Units |
| --- | --- | --- |
| $C$ | 1 | $\mu F/cm^2$ |
| $g_{Na}$ | 50 | $mS/cm^2$ |
| $g_K$ | 5 | $mS/cm^2$ |
| $g_L$ | 0.1 | $mS/cm^2$ |
| $g_{LD}$ | 0.01 | $mS/cm^2$ |
| $g_P$ | 0.06 | $mS/cm^2$ |
| $g_{AHP}$ | 0.8 | $mS/cm^2$ |
| $g_{Ca}$ | 0.25 | $mS/cm^2$ |
| $g_d$ | 0.3 | $mS/cm^2$ |
| $V_K$ | -110 | $mV$ |
| $V_L$ | -65 | $mV$ |
| $V_{LD}$ | -67 | $mV$ |
| $V_{Ca}$ | 120 | $mV$ |
| $V_{Na}$ | 40 | $mV$ |

## References

Aghajanian GK, Cedarbaum JM & Wang RY. (1977). Evidence for norepinephrine-mediated collateral inhibition of locus coeruleus neurons. Brain Res 136, 570–577.

Aghajanian GK & VanderMaelen CP. (1982). alpha 2-adrenoceptor-mediated hyperpolarization of locus coeruleus neurons: intracellular studies in vivo. Science 215, 1394–1396.

Alvarez VA, Chow CC, Van Bockstaele EJ & Williams JT. (2002). Frequency-dependent synchrony in locus ceruleus: role of electrotonic coupling. Proc Natl Acad Sci U S A 99, 4032–4036.

Andrade R & Aghajanian GK. (1984). Intrinsic regulation of locus coeruleus neurons: electrophysiological evidence indicating a predominant role for autoinhibition. Brain Res 310, 401–406.

Aston-Jones G & Bloom FE. (1981). Activity of norepinephrine-containing locus coeruleus neurons in behaving rats anticipates fluctuations in the sleep-waking cycle. J Neurosci 1, 876–886.

Aston-Jones G & Cohen JD. (2005). Adaptive gain and the role of the locus coeruleus-norepinephrine system in optimal performance. J Comp Neurol 493, 99–110.

Aston-Jones G & Waterhouse B. (2016). Locus coeruleus: From global projection system to adaptive regulation of behavior. Brain Res 1645, 75–78.

Baral S, Hosseini H, More K, Fabrin TMC, Braun J & Prigge M. (2022). Spike-Dependent Dynamic Partitioning of the Locus Coeruleus Network through Noradrenergic Volume Release in a Simulation of the Nucleus Core. Brain Sci 12.

Benarroch EE. (2009). The locus ceruleus norepinephrine system: functional organization and potential clinical significance. Neurology 73, 1699–1704.

Berridge CW & Waterhouse BD. (2003). The locus coeruleus-noradrenergic system: modulation of behavioral state and state-dependent cognitive processes. Brain Res Brain Res Rev 42, 33–84.

Breton-Provencher V, Drummond GT, Feng J, Li Y & Sur M. (2022). Spatiotemporal dynamics of noradrenaline during learned behaviour. Nature 606, 732–738.

Breton-Provencher V, Drummond GT & Sur M. (2021). Locus Coeruleus Norepinephrine in Learned Behavior: Anatomical Modularity and Spatiotemporal Integration in Targets. Front Neural Circuits 15, 638007.

Breton-Provencher V & Sur M. (2019). Active control of arousal by a locus coeruleus GABAergic circuit. Nat Neurosci 22, 218–228.

Callado LF & Stamford JA. (1999). Alpha2A-but not alpha2B/C-adrenoceptors modulate noradrenaline release in rat locus coeruleus: voltammetric data. Eur J Pharmacol 366, 35–39.

Carter ME, Yizhar O, Chikahisa S, Nguyen H, Adamantidis A, Nishino S, Deisseroth K & de Lecea L. (2010). Tuning arousal with optogenetic modulation of locus coeruleus neurons. Nat Neurosci 13, 1526–1533.

Cedarbaum JM & Aghajanian GK. (1977). Catecholamine receptors on locus coeruleus neurons: pharmacological characterization. Eur J Pharmacol 44, 375–385.

Cedarbaum JM & Aghajanian GK. (1978). Activation of locus coeruleus neurons by peripheral stimuli: modulation by a collateral inhibitory mechanism. Life Sci 23, 1383–1392.

Chandler DJ, Gao WJ & Waterhouse BD. (2014). Heterogeneous organization of the locus coeruleus projections to prefrontal and motor cortices. Proc Natl Acad Sci U S A 111, 6816–6821.

Chandler DJ, Jensen P, McCall JG, Pickering AE, Schwarz LA & Totah NK. (2019). Redefining Noradrenergic Neuromodulation of Behavior: Impacts of a Modular Locus Coeruleus Architecture. J Neurosci 39, 8239–8249.

Cherubini E, North RA & Williams JT. (1988). Synaptic potentials in rat locus coeruleus neurones. J Physiol 406, 431–442.

Christie MJ, Williams JT & North RA. (1989). Electrical coupling synchronizes subthreshold activity in locus coeruleus neurons in vitro from neonatal rats. J Neurosci 9, 3584–3589.

Courtney NA & Ford CP. (2014). The timing of dopamine- and noradrenaline-mediated transmission reflects underlying differences in the extent of spillover and pooling. J Neurosci 34, 7645–7656.

Dahlstroem A & Fuxe K. (1964). Evidence for the Existence of Monoamine-Containing Neurons in the Central Nervous System. I. Demonstration of Monoamines in the Cell Bodies of Brain Stem Neurons. Acta Physiol Scand Suppl, SUPPL 232:231–255.

Dvorkin R & Shea SD. (2022). Precise and Pervasive Phasic Bursting in Locus Coeruleus during Maternal Behavior in Mice. J Neurosci 42, 2986–2999.

Egan TM, Henderson G, North RA & Williams JT. (1983). Noradrenaline-mediated synaptic inhibition in rat locus coeruleus neurones. J Physiol 345, 477–488.

Ennis M & Aston-Jones G. (1986). Evidence for self- and neighbor-mediated postactivation inhibition of locus coeruleus neurons. Brain Res 374, 299–305.

Ermentrout GB & Terman DH. (2010). Mathematical Foundations of Neuroscience, 10.1007/978-0-387-87708-2. Springer, New York.

Favre-Bulle IA, Vanwalleghem G, Taylor MA, Rubinsztein-Dunlop H & Scott EK. (2018). Cellular-Resolution Imaging of Vestibular Processing across the Larval Zebrafish Brain. Curr Biol 28, 3711–3722 e3713.

Feng J, Zhang C, Lischinsky JE, Jing M, Zhou J, Wang H, Zhang Y, Dong A, Wu Z, Wu H, Chen W, Zhang P, Zou J, Hires SA, Zhu JJ, Cui G, Lin D, Du J & Li Y. (2019). A Genetically Encoded Fluorescent Sensor for Rapid and Specific In Vivo Detection of Norepinephrine. Neuron 102, 745–761 e748.

Foote SL, Bloom FE & Aston-Jones G. (1983). Nucleus locus ceruleus: new evidence of anatomical and physiological specificity. Physiol Rev 63, 844–914.

Gong R, Xu S, Hermundstad A, Yu Y & Sternson SM. (2020). Hindbrain Double-Negative Feedback Mediates Palatability-Guided Food and Water Consumption. Cell 182, 1589–1605 e1522.

Grandoso L, Pineda J & Ugedo L. (2004). Comparative study of the effects of desipramine and reboxetine on locus coeruleus neurons in rat brain slices. Neuropharmacology 46, 815–823.

Groves PM & Wilson CJ. (1980). Monoaminergic presynaptic axons and dendrites in rat locus coeruleus seen in reconstructions of serial sections. J Comp Neurol 193, 853–862.

Hayat H, Regev N, Matosevich N, Sales AC, Paredes-Rodriguez E, Krom AJ, Bergman L, Li Y, Lavigne M, Kremer EJ, Yizhar O, Pickering AE & Nir Y. (2020). Locus-coeruleus norepinephrine activity mediates sensory-evoked awakenings from sleep. Science Advances 6, eaaz4232.

Hickey L, Li Y, Fyson SJ, Watson TC, Perrins R, Hewinson J, Teschemacher AG, Furue H, Lumb BM & Pickering AE. (2014). Optoactivation of locus ceruleus neurons evokes bidirectional changes in thermal nociception in rats. J Neurosci 34, 4148–4160.

Hirschberg S, Li Y, Randall A, Kremer EJ & Pickering AE. (2017). Functional dichotomy in spinal- vs prefrontal-projecting locus coeruleus modules splits descending noradrenergic analgesia from ascending aversion and anxiety in rats. eLife 6.

Howorth P, Thornton S, O’Brien V, Smith W, Nikiforova N, Teschemacher A & Pickering A. (2009a). Retrograde Viral Vector-Mediated Inhibition of Pontospinal Noradrenergic Neurons Causes Hyperalgesia in Rats. Journal of Neuroscience 29, 12855–12864.

Howorth PW, Teschemacher AG & Pickering AE. (2009b). Retrograde adenoviral vector targeting of nociresponsive pontospinal noradrenergic neurons in the rat in vivo. J Comp Neurol 512, 141–157.

Huang HP, Wang SR, Yao W, Zhang C, Zhou Y, Chen XW, Zhang B, Xiong W, Wang LY, Zheng LH, Landry M, Hokfelt T, Xu ZQ & Zhou Z. (2007). Long latency of evoked quantal transmitter release from somata of locus coeruleus neurons in rat pontine slices. Proc Natl Acad Sci U S A 104, 1401–1406.

Ishimatsu M & Williams JT. (1996). Synchronous activity in locus coeruleus results from dendritic interactions in pericoerulear regions. J Neurosci 16, 5196–5204.

Kebschull JM, Garcia da Silva P, Reid AP, Peikon ID, Albeanu DF & Zador AM. (2016). High-Throughput Mapping of Single-Neuron Projections by Sequencing of Barcoded RNA. Neuron 91, 975–987.

Kjaerby C, Andersen M, Hauglund N, Untiet V, Dall C, Sigurdsson B, Ding F, Feng J, Li Y, Weikop P, Hirase H & Nedergaard M. (2022). Memory-enhancing properties of sleep depend on the oscillatory amplitude of norepinephrine. Nat Neurosci 25, 1059–1070.

Kuo CC, Hsieh JC, Tsai HC, Kuo YS, Yau HJ, Chen CC, Chen RF, Yang HW & Min MY. (2020). Inhibitory interneurons regulate phasic activity of noradrenergic neurons in the mouse locus coeruleus and functional implications. J Physiol 598, 4003–4029.

Li Y, Hickey L, Perrins R, Werlen E, Patel AA, Hirschberg S, Jones MW, Salinas S, Kremer EJ & Pickering AE. (2016). Retrograde optogenetic characterization of the pontospinal module of the locus coeruleus with a canine adenoviral vector. Brain Res 1641, 274–290.

Llorca-Torralba M, Borges G, Neto F, Mico JA & Berrocoso E. (2016). Noradrenergic Locus Coeruleus pathways in pain modulation. Neuroscience 338, 93–113.

Loughlin SE, Foote SL & Bloom FE. (1986). Efferent projections of nucleus locus coeruleus: topographic organization of cells of origin demonstrated by three-dimensional reconstruction. Neuroscience 18, 291–306.

Magnus CJ, Lee PH, Atasoy D, Su HH, Looger LL & Sternson SM. (2011). Chemical and genetic engineering of selective ion channel-ligand interactions. Science 333, 1292–1296.

Matosevich N & Nir Y. (2021). Noradrenaline: Sleep on it. Curr Biol 31, R1477–R1479.

Morici JF & Girardeau G. (2022). Cortical norepinephrine GRABs a seat at the sleep table. Nat Neurosci 25, 978–980.

Nadkarni S, Bartol TM, Sejnowski TJ & Levine H. (2010). Modelling vesicular release at hippocampal synapses. PLoS computational biology 6, e1000983.

Osorio-Forero A, Cardis R, Vantomme G, Guillaume-Gentil A, Katsioudi G, Devenoges C, Fernandez LMJ & Luthi A. (2021). Noradrenergic circuit control of non-REM sleep substates. Curr Biol 31, 5009–5023 e5007.

Patel M & Joshi B. (2015). Modeling the evolving oscillatory dynamics of the rat locus coeruleus through early infancy. Brain Res 1618, 181–193.

Plummer NW, Scappini EL, Smith KG, Tucker CJ & Jensen P. (2017). Two Subpopulations of Noradrenergic Neurons in the Locus Coeruleus Complex Distinguished by Expression of the Dorsal Neural Tube Marker Pax7. Front Neuroanat 11, 60.

Poe GR, Foote S, Eschenko O, Johansen JP, Bouret S, Aston-Jones G, Harley CW, Manahan-Vaughan D, Weinshenker D, Valentino R, Berridge C, Chandler DJ, Waterhouse B & Sara SJ. (2020). Locus coeruleus: a new look at the blue spot. Nat Rev Neurosci 10.1038/s41583-020-0360-9.

Pudovkina OL, Kawahara Y, de Vries J & Westerink BH. (2001). The release of noradrenaline in the locus coeruleus and prefrontal cortex studied with dual-probe microdialysis. Brain Res 906, 38–45.

Romano SA, Perez-Schuster V, Jouary A, Boulanger-Weill J, Candeo A, Pietri T & Sumbre G. (2017). An integrated calcium imaging processing toolbox for the analysis of neuronal population dynamics. PLoS computational biology 13, e1005526.

Sales AC, Friston KJ, Jones MW, Pickering AE & Moran RJ. (2019). Locus Coeruleus tracking of prediction errors optimises cognitive flexibility: An Active Inference model. PLoS computational biology 15, e1006267.

Samuels ER & Szabadi E. (2008). Functional neuroanatomy of the noradrenergic locus coeruleus: its roles in the regulation of arousal and autonomic function part II: physiological and pharmacological manipulations and pathological alterations of locus coeruleus activity in humans. Current neuropharmacology 6, 254–285.

Sara SJ. (2009). The locus coeruleus and noradrenergic modulation of cognition. Nat Rev Neurosci 10, 211–223.

Savitzky A & Golay MJE. (1964). Smoothing and Differentiation of Data by Simplified Least Squares Procedures. Analytical Chemistry 36, 1627–1639.

Schindelin J, Arganda-Carreras I, Frise E, Kaynig V, Longair M, Pietzsch T, Preibisch S, Rueden C, Saalfeld S, Schmid B, Tinevez JY, White DJ, Hartenstein V, Eliceiri K, Tomancak P & Cardona A. (2012). Fiji: an open-source platform for biological-image analysis. Nat Methods 9, 676-682.

Schwarz LA, Miyamichi K, Gao XJ, Beier KT, Weissbourd B, DeLoach KE, Ren J, Ibanes S, Malenka RC, Kremer EJ & Luo L. (2015). Viral-genetic tracing of the input-output organization of a central noradrenaline circuit. Nature 10.1038/nature14600.

Sharma Y, Xu T, Graf WM, Fobbs A, Sherwood CC, Hof PR, Allman JM & Manaye KF. (2010). Comparative anatomy of the locus coeruleus in humans and nonhuman primates. J Comp Neurol 518, 963–971.

Shimizu N & Imamoto K. (1970). Fine structure of the locus coeruleus in the rat. Arch Histol Jpn 31, 229–246.

Singewald N, Schneider C, Pfitscher A & Philippu A. (1994). In vivo release of catecholamines in the locus coeruleus. Naunyn Schmiedebergs Arch Pharmacol 350, 339–345.

Sudhof TC. (2012). Calcium control of neurotransmitter release. Cold Spring Harb Perspect Biol 4, a011353.

Swanson LW. (1976). The locus coeruleus: a cytoarchitectonic, Golgi and immunohistochemical study in the albino rat. Brain Res 110, 39–56.

Tavares A, Handy DE, Bogdanova NN, Rosene DL & Gavras H. (1996). Localization of alpha 2A- and alpha 2B-adrenergic receptor subtypes in brain. Hypertension 27, 449–455.

Totah NK, Neves RM, Panzeri S, Logothetis NK & Eschenko O. (2018). The Locus Coeruleus Is a Complex and Differentiated Neuromodulatory System. Neuron 99, 1055–1068 e1056.

Tucker WC & Chapman ER. (2002). Role of synaptotagmin in Ca2+-triggered exocytosis. Biochem J 366, 1–13.

Uematsu A, Tan BZ, Ycu EA, Cuevas JS, Koivumaa J, Junyent F, Kremer EJ, Witten IB, Deisseroth K & Johansen JP. (2017). Modular organization of the brainstem noradrenaline system coordinates opposing learning states. Nat Neurosci 10.1038/nn.4642.

Vanwalleghem G, Constantin L & Scott EK. (2020). Calcium Imaging and the Curse of Negativity. Front Neural Circuits 14, 607391.

Virtanen P, Gommers R, Oliphant TE, Haberland M, Reddy T, Cournapeau D, Burovski E, Peterson P, Weckesser W, Bright J, van der Walt SJ, Brett M, Wilson J, Millman KJ, Mayorov N, Nelson ARJ, Jones E, Kern R, Larson E, Carey CJ, Polat I, Feng Y, Moore EW, VanderPlas J, Laxalde D, Perktold J, Cimrman R, Henriksen I, Quintero EA, Harris CR, Archibald AM, Ribeiro AH, Pedregosa F, van Mulbregt P & SciPy C. (2020). SciPy 1.0: fundamental algorithms for scientific computing in Python. Nat Methods 17, 261–272.

Wagner-Altendorf TA, Fischer B & Roeper J. (2019). Axonal projection-specific differences in somatodendritic alpha2 autoreceptor function in locus coeruleus neurons. Eur J Neurosci 10.1111/ejn.14553.

Weinshenker D. (2018). Long Road to Ruin: Noradrenergic Dysfunction in Neurodegenerative Disease. Trends Neurosci 41, 211–223.

Williams JT, Henderson G & North RA. (1985). Characterization of alpha 2-adrenoceptors which increase potassium conductance in rat locus coeruleus neurones. Neuroscience 14, 95–101.

Williams JT & Marshall KC. (1987). Membrane properties and adrenergic responses in locus coeruleus neurons of young rats. J Neurosci 7, 3687–3694.

Williams JT & North RA. (1985). Catecholamine inhibition of calcium action potentials in rat locus coeruleus neurones. Neuroscience 14, 103–109.

Williams JT, North RA, Shefner SA, Nishi S & Egan TM. (1984). Membrane properties of rat locus coeruleus neurones. Neuroscience 13, 137–156.

Zoli M, Torri C, Ferrari R, Jansson A, Zini I, Fuxe K & Agnati LF. (1998). The emergence of the volume transmission concept. Brain Res Brain Res Rev 26, 136–147.

